# Impact of brain parcellation on prediction performance in models of cognition and demographics

**DOI:** 10.1101/2023.02.10.528041

**Authors:** Marta Czime Litwińczuk, Nils Muhlert, Nelson Trujillo-Barreto, Anna Woollams

## Abstract

Brain connectivity analysis begins with the selection of a parcellation scheme that will define brain regions as nodes of a network whose connections will be studied. Brain connectivity has already been used in predictive modelling of cognition, but it remains unclear if the resolution of the parcellation used can systematically impact the predictive model performance. In this work, structural, functional and combined connectivity were each defined with 5 different parcellation schemes. The resolution and modality of the parcellation schemes were varied. Each connectivity defined with each parcellation was used to predict individual differences in age, education, sex, Executive Function, Self-regulation, Language, Encoding and Sequence Processing. It was found that low-resolution functional parcellation consistently performed above chance at producing generalisable models of both demographics and cognition. However, no single parcellation scheme proved superior at predictive modelling across all cognitive domains and demographics. In addition, although parcellation schemes impacted the global organisation of each connectivity type, this difference could not account for the out-of-sample prediction performance of the models. Taken together, these findings demonstrate that while high-resolution parcellations may be beneficial for modelling specific individual differences, partial voluming of signals produced by higher resolution of parcellation likely disrupts model generalisability.

## INTRODUCTION

Neuroimaging research has demonstrated that adaptive behaviour relies not only on localised activation in the brain but also on effective coordination of activity across remote neuronal populations (Alvarez & Squire, 1994; Seeley et al., 2007). To understand how information is exchanged in the human brain, research investigates the white matter connections between regions (structural connectivity, SC) and the statistical associations between their activity (functional connectivity, FC)(Park & Friston, 2013). SC and FC have been related to healthy cognitive function throughout the lifespan (Salami et al., 2014), whereas their disruption has been demonstrated to characterise many psychiatric, developmental and clinical diagnostics (de Kwaasteniet et al., 2013; Guye et al., 2010; Hahn et al., 2013; Keller et al., 2007; van den Heuvel & Fornito, 2014). Consequently, the study of the relationship between brain connectivity and cognition has become increasingly popular (Farahani et al., 2019; Hoppenbrouwers et al., 2014; Manca et al., 2018).

One of the most central questions in the analysis of brain connectivity is how to define the brain regions (i.e. brain parcellation) used for connectivity analysis (Eickhoff et al., 2018; Yao et al., 2015). In the analysis of brain connectivity, the brain is idealised (modelled) as a set of interconnected nodes each representing a brain region that is assumed to constitute a coherent unit of activity. The connectivity pattern between regions is then studied and often used to explain and predict cognitive function. Therefore, brain parcellation is a key step of any connectivity analysis. Nearly two dozen different brain parcellations are now available (Arslan et al., 2018; Dickie et al., 2017; Eickhoff et al., 2018; Lawrence et al., 2021). Each parcellation has been defined using different neuroimaging data and parcel generation algorithms based on different parcellation criteria. Consequently, each parcellation offers different node locations, sizes and shapes therefore leading to different representations of the activity units used for analysis. However, clinical and cognitive neuroimaging research does not yet have a standard or most common parcellation. In addition, it is unlikely that a ‘one size fits all’ standard will emerge because every parcellation method has been defined to highlight specific properties of the brain (Eickhoff et al., 2018). To illustrate, we may choose to parcellate the brain based on histological boundaries (Amunts & Zilles, 2015; Brodmann, 1909) or clustering of a group of similarly active voxels (Cohen et al., 2008). Such choices appear to impact consistency of research findings (Dhamala et al., 2021; Mellema et al., 2020; Ota et al., 2014; Pervaiz et al., 2020).

Despite the lack of an accepted standard, parcellation choice is an important decision in network analysis and it carries profound consequences for interpretation of results and comparison across different studies. It has been empirically demonstrated that parcellation can impact the properties of the estimated brain networks. For example, the parcellation used can impact the small world coefficient of SC and FC, which refers to the network’s tendency to segregate into clusters of strongly connected nodes relative to its tendency to produce short paths between any pair of nodes (Fornito, 2010; Wang et al., 2009; Zalesky et al., 2010). In addition, the ability to produce highly connected nodes (aka degree distribution) can also be affected by the specific parcellation used (Fornito, 2010; Wang et al., 2009; Zalesky et al., 2010). Similarly, the choice of parcellation resolution has also been demonstrated to impact the similarities between structural and functional connectivity (Akiki & Abdallah, 2019; Ashourvan et al., 2019; Honey et al., 2009).

Parcellation choice alone has been demonstrated to impact the explanatory and predictive power of neuroimaging models of cognition in healthy and clinical populations (Dhamala et al., 2021; Mellema et al., 2020; Ota et al., 2014; Pervaiz et al., 2020). In particular, some evidence suggests that high-resolution parcellations may benefit the predictive power of connectivity models of cognition (Mellema et al., 2020; Pervaiz et al., 2020). The difference in predictive power of various brain parcellations may be related to the properties of the networks that become more prominent with specific parcellations. For example, increased small world characteristics of FC have been related to high intelligence (Hilger et al., 2017; Langer et al., 2012). Therefore, if one brain parcellation tends to generate networks with increased specific properties, then this parcellation may influence the accuracy of model predictions. However, it is also possible that difference in predictive power of specific models of cognition is related to whether specific parcellations generate accurate connectivity profiles. For example, it is likely that a functionally-defined parcellation is less prone to partial voluming of functional signals as compared to structurally-defined parcellation. In contrast, a structurally-defined parcellation that considers anatomical boundaries of regions may be better suited to define SC. In support of this proposal, Dhamala et al. (2021) have found a trend that SC defined with structural parcellation yielded more effective models of crystallised cognition than SC defined with activation-based parcellation.

However, no work has yet been conducted to assess if the parcellation method systematically influences network architecture and the explanatory and/or predictive power of models of cognition. The present work investigated this question by defining SC and FC with a variety of parcellations: the resting state activity-defined parcellation with either 93, 184 and 278 parcels (Shen et al., 2013), the structurally-defined parcellation (Rolls et al., 2020) and a hybrid parcellation that considers both structural and functional information (Xia et al., 2013). Our work then produced connectivity-based predictive models of cognition and used graph theory to assess the global differences in network properties that may explain model differences in model performance. Doing so, we tested the hypotheses that (1) SC-based models of demographics and cognition will be more effective when connectivity is defined with a structural parcellation (Rolls et al., 2020) than with a parcellation based on a functional activation (Shen et al., 2013), (2) FC-based models of demographics and cognition will be more effective when connectivity is defined with a parcellation based on brain function (Shen et al., 2013) than with a parcellation based on structure, (3) use of high-resolution parcellation will improve explanatory and predictive power of models of demographics and cognition regardless of connectivity type, (4) differences in model explanatory and predictive power will systematically map with differences in network organisation, as measured by graph theory.

## METHODS

### Participants

Neuroimaging and cognitive data were obtained for 250 unrelated subjects from the 1200-subject release of the Human Connectome Project (HCP). For consistent treatment of behavioural and neuroimaging subjects’ data selection, one subject was excluded from the neuroimaging analysis due to incomplete behavioural data. The sample consisted of 138 females and 111 males in the age range of 22 and 36 years.

### Measures of cognition

The present work used the same measures of cognition as our previous work (Litwińczuk et al., 2022). Analysed tasks included: Picture Sequence Memory, Dimensional Change Card Sort, Flanker Inhibitory Control and Attention Task, Penn Progressive Matrices, Oral Reading Recognition, Picture Vocabulary, Pattern Comparison Processing Speed, Delay Discounting, Variable Short Penn Line Orientation Test, Short Penn Continuous Performance Test, Penn Word Memory Test, and List Sorting. These assessments were obtained from the Blueprint for Neuroscience Research–funded NIH Toolbox for Assessment of Neurological and Behavioral function (http://www.nihtoolbox.org) and tasks from the Penn computerized neurocognitive battery (Gur et al., 2010). Principal Component Analysis (PCA) with VARIMAX rotation was then used as a feature extraction method and applied to the behavioural dataset. The extracted PCA rotated components reflected specific latent cognitive domains, interpreted as Executive Function, Self-regulation, Language, Encoding and Sequence Processing. The present work uses the PCA scores obtained previously for each cognitive domain.

### Minimally processed neuroimaging data

The HCP provides minimally processed neuroimaging data that was used here, the data acquisition and processing pipeline has been discussed in detail by (Glasser et al., 2013). All neuroimaging data were collected with a 3T Siemens “Connectome Skyra” scanner that uses the Siemens 32-channel RF receive head coil and with an SC72 gradient insert (Ugurbil et al., 2013). Here, we utilised Version 3 of the minimal processing pipeline implemented with FSL 5.0.6 (Jenkinson et al., 2012) and FreeSurfer 5.3.0-HCP (Dale et al., 1999).

T1 weighted MR images were acquired with a 3D MPRAGE sequence (TR = 2400 ms, TE = 2.14, TI = 1000 ms, flip angle = 8°, FOV = 224 by 224 mm, voxel size = 0.7 mm isotropic). Rs-fMRI data was collected using the gradient-echo EPI (TR = 720 ms, TE = 33.1 ms, flip angle = 52°, FOV = 208 by 180 mm, 70 slices, thickness = 2.0 mm, size = 2.0 mm isotropic). Scans were collected in two sessions, each lasting approximately 15 minutes. The rs-fMRI data were collected both in left-to-right and right-to-left directions. In addition, in the original data, spin echo phase reversed images were acquired for registration with T1 images and the spin echo field maps were acquired for bias field correction. Diffusion-weighted MR images were acquired with spin-echo EPI sequence (TR = 5520 ms, TE = 89.5 ms, flip angle = 78°, refocusing flip angle = 160°, FOV = 210 by 180 mm, 111 slices, thickness = 1.25 mm, size = 1.25 mm isotropic). Each gradient consisted of 90 diffusion weighting directions plus 6 b=0. There were 3 diffusion-weighed shells of b=1000, 2000, and 3000 s/mm^2^ . SENSE1 multi-channel image reconstruction was used (Sotiropoulos et al., 2013).

### Additional processing of neuroimaging data

The Neuroimaging data were further processed following the same processing pipeline as in our previous work (Litwińczuk et al., 2022). This pipeline is summarised for completeness in the next two sub-sections. Both SC and FC were defined with 5 parcellation schemes that varied in their modality and resolution. For clarity, throughout this work the parcellation name will be assisted with the number of parcels:

1. Low-resolution resting-state functional parcellation composed of 93 parcels (Shen et al., 2013)
2. Structural parcellation known as Automated Anatomical Labelling (AAL3) composed of 166 parcels (Rolls et al., 2020)
3. Moderate-resolution resting-state functional parcellation composed of 184 parcels (Shen et al., 2013)
4. Hybrid structural and functional parcellation known as Brainnetome composed of 246 parcels (Xia et al., 2013).
5. High-resolution resting-state functional parcellation composed of 278 parcels (Shen et al., 2013)

#### Structural data and structural connectivity calculation

The diffusion data were further analysed using the BEDPOSTX procedure in FSL, which runs Markov Chain Monte Carlo sampling to estimate probability distributions on diffusion parameters at each voxel. This information was used in the FDT module of FSL to run ROI-to-ROI probabilistic tractography with ProbtrackX. Tractography was run between parcels in each of the 5 parcellations.

During tractography, 5000 streamlines were initiated from each voxel with a step length of 0.5 mm (Behrens et al., 2007; Behrens et al., 2003; Jenkinson et al., 2012). Streamlines were constrained with a curvature threshold of 0.2, a maximum of 2000 steps per streamline and a volume fraction threshold of subsidiary fiber orientations of 0.01. An SC matrix between regions was constructed by first counting the number of streamlines originating from a seed region *i* that reached a target region *j* (*S_ij_*). These counts are asymmetric since the count of streamlines from region *i* to *j* is not necessarily equal to the count of streamlines from region *j* to *i* (*S_ij_* ≠ *S_ji_*), but they are highly correlated for all subjects (lowest Pearson’s Correlation was 0.76, p < 0.001). Based on these counts, the weight *W_ij_* (entries of the SC matrix) between any two pairs of regions *i* and *j* was defined as the ratio of the total streamline counts in both directions (*S_ij_* + *S_ji_*), to the maximum possible number of streamlines that can be shared between the two regions, which is (*N_i_* + *N_j_*) * 5000 (where *N_i_* and *N_j_* are the numbers of seed voxels in regions *i* and *j*, respectively):,

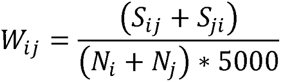

Similar to previous studies, the weight *W_ij_* can be interpreted as capturing the connection density (number of streamlines per unit surface) between nodes *i* and *j*, which accounts for possible bias due to the different sizes of the seed regions (Friston et al., 2008; Ingalhalikar et al., 2013). Note that the SC matrix defined based on these weights is symmetric because swapping around the regions’ indices does not change the result; and it is also normalised between 0 and 1, because the maximum value of the numerator can only be reached when all streamlines originating from each of region reach the other region, so that *S_ij_* = *N_i_* * 5000 and *S_ji_* = *N_j_* * 5000, which gives *W_ij_* = 1. Evidence suggests that structural connectivity is most sensitive to individual differences with moderate-to-high thresholding (Buchanan et al., 2020) and produces the least false positive and negative results (de Reus & van den Heuvel, 2013), therefore an 80% proportional threshold was applied.

#### Functional data and functional connectivity calculation

The minimally processed rs-fMRI data was used to compute FC based on pair-wise correlations (Glasser et al., 2013). Next, the following steps were taken to further process data using the CONN Toolbox (Whitfield-Gabrieli & Nieto-Castanon, 2012) with the use of the standard FC processing pipeline (Nieto-Castanon, 2020). Briefly, images were realigned, a slice-timing corrected, and outlier detection of functional images for scrubbing was performed with Artefact Detection Tools (ART, https://www.nitrc.org/projects/artifact_detect/). Grey matter, white matter, cerebrospinal fluid and non-brain tissues were then segmented. Images were normalized and smoothed with a 6 mm Full Width at Half Maximum Gaussian kernel. Next, the data was denoised with default Conn denoising options using the anatomical component-based noise correction procedure (Behzadi et al., 2007). This procedure removes artefactual components from the data, including noise components from cerebral white matter and cerebrospinal areas, subject-motion parameters (Friston et al., 1996), identified outlier scans (Power et al., 2014), and constant and first-order linear session effects (Whitfield-Gabrieli & Nieto-Castanon, 2012). Then standard denoising steps were applied including scrubbing, motion regression and application of a high pass filter (0.01 Hz cut-off), and a low pass filter (0.10 Hz cut-off).

FC analysis was performed based on the same 5 parcellations. The average blood oxygenation level-dependent signal in each parcel was obtained and the pairwise (parcel-to-parcel) correlation of the averaged signals was calculated. Since the CONN toolbox produces Fisher’s Z-scores (Fisher, 1915), a hyperbolic tangent function was used to reverse Fisher’s transformation and obtain original correlation values ranging between −1 and 1. Negative correlations were transformed to positive by taking their absolute values and a proportional 80% FC threshold was then applied (Garrison et al., 2015; van den Heuvel et al., 2017).

### Model construction

Predictive models of 5 cognitive domains and 3 demographic variables were separately constructed using FC, SC or a combination of SC and FC (referred to as combined connectivity (CC)) as predictors. Models also differed in the brain parcellation scheme used to define the network nodes for connectivity calculations. This led to a total of 120 models to be analysed ((5 cognitive domains + 3 demographic characteristics) * 3 connectivity types * 5 parcellation schemes). Prior to model estimation, for each cognitive domain, the confounding effect of age, gender and education was regressed out of the response variable. Meanwhile, for each demographic, the remaining demographics were included in the main model as covariates of no interest. Then, all models were estimated using a Principal Component Regression (PCR) approach with elastic net regularisation of the regression coefficients in latent space (EN-PCR). The overall estimation procedure consisted of the following steps:

1. PCA decomposition was used to orthogonalise the predictors’ (connectivity) matrix
2. A regression model in latent (PCA) space was fitted with elastic net regularisation for the regression coefficients
3. Regression coefficients obtained in PCA space were projected back to the original connectome space and used to produce predicted responses.

To tune the Elastic Net regularisation hyper-parameters alpha and lambda and evaluate the out-of-sample model performance, a Bootstrap Bias Corrected Cross-Validation (BBC-CV) approach was used (Tsamardinos et al., 2018). In brief, the BBC-CV consisted in a repeated (50 times) 5-fold cross-validation with hyperparameter tuning (CVT), followed by a Bootstrap bias correction procedure (5000 bootstrap samples). The later Bootstrap step accounts for optimistic biases in the estimation of model performance introduced by using the same data for both hyperparameter tuning and model evaluation in the CVT step (Tsamardinos et al., 2018). The explained variation (coefficient of determination) between the predicted and observed responses was used as the statistic for both hyperparameter tuning and out-of-sample performance evaluation of the model (Poldrack et al., 2020). The resultant population of bootstrapped statistics was used to produce mean performance estimates of the EN-PCR learning strategy and corresponding confidence intervals. For a similar application of the BBC-CV method in the context of predictive modelling of cognition based on connectivity we refer the authors to our previous paper (Litwińczuk et al., 2022).

Finally, a permutation test was used to assess how likely it is to get the observed models’ performance by chance. Specifically, the out-of-sample predictions produced by the BBC-CVT (249 predictions) were permuted (sampled without replacement) 5,000 times and the models’ performance statistics (coefficient of determination) were estimated for each permutation. This null distribution was then used to assess the observed model performance statistics in the non-permuted data. That is, the p-value for testing the significance of models’ performance was determined by computing the proportion of resampled statistics at least as high as or greater than the observed statistics.

### Model comparison

All following analysis was done separately for each cognitive domain and connectivity type, and the analysis compared the effects of parcellation choice.

To compare the models’ whole-sample explanatory power, the Akaike Information Criterion (AIC) for each model was obtained. The AIC balances the goodness-of-fit of the model (model accuracy) against its complexity (the number of estimable parameters of the model) so that a reduced AIC is associated with improved model quality. Given any two models *M*_l_ and *M*_2_, a positive difference (ΔAIC = AIC(*M*_l_)-AIC(*M*_2_)) is interpreted as weak (1-2 units), strong (2-4 units), considerably strong (4-7 units) and decisive (>10) evidence against evidence against *M*_l_ (Burnham & Anderson, 2016). We used the AIC difference to compare the explanatory power of alternative models of the same response variable (cognitive or demographic) with the same connectivity modality as predictor (SC, FC or CC). The models under comparison only differed in the type of parcellation used to compute the connectivity predictors. Consequently, if two parcellations had similar explanatory power but one was coarser than the other, the coarser parcellation was favoured due to requiring fewer parameters (number of connectivity predictors) to model the same response data.

To compare the models’ out-of-sample predictive performance, the models’ out-of-sample predictions (generated during repeated BBC-CVT) were used to obtain a bootstrapped (5000 samples) estimates of the models’ performance statistics (coefficient of determination). Additionally, the non-parametric Wilcoxon rank sum test for equal medians was used to assess the significance of differences in performance between different models. These comparisons were only done for models which performed better than chance.

### Graph theory measures

Graph theoretic measures were calculated based on the weighted, undirected SC and FC matrices of every subject, using The Brain Connectivity Toolbox (http://www.brain-connectivity-toolbox.net). Measures of global network organisation included small-world propensity, global efficiency, assortativity, modularity statistic, transitivity, and coreness statistic resulting from core-periphery partition. Small world propensity requires computation of the clustering coefficient and the shortest path length of the network (Muldoon et al., 2016). The clustering coefficient was obtained using Onnela’s algorithm (Muldoon et al., 2016; Onnela et al., 2005) whereas the shortest path length was obtained using the Floyd–Warshall Algorithm applied to the weighted graph obtained from the inverse of each connectivity matrix. Modularity statistic requires computation of network modules, which were defined with Newman’s algorithm (Newman, 2006).

To study the impact of the parcellation on global network organisation, a series of paired t-tests were used to test for significant differences between the same global graph theoretic measure computed for two different parcellation schemes. To explore the impact of global network architecture yielded by each parcellation on predictive modelling, regression models were fitted to each observed cognitive domain using graph theory. Graph theory models were fitted and compared in the same manner as for connectivity but without elastic-net regularisation.

## RESULTS

### Connectivity

#### Predictive modelling of demographics

##### AIC-based model comparison

AIC values demonstrated that across all connectivity modalities and parcellations, age was best modelled with SC defined either with AAL3 (166) or Shen (93) parcellations (Figure 1). The difference between AIC values of SC models defined with either AAL3 (166) or Shen (93) was small. For SC, there was evidence in favour of both AAL3 (166) and Shen (93) parcellations relative to Shen (184) and Shen (278), and Brainnetome (246). For FC, there was substantial evidence in favour of Shen (93) parcellation to AAL3 (166). For CC, there was substantial evidence in favour of Shen (93) parcellation above to Shen (184).

**Figure 1.**
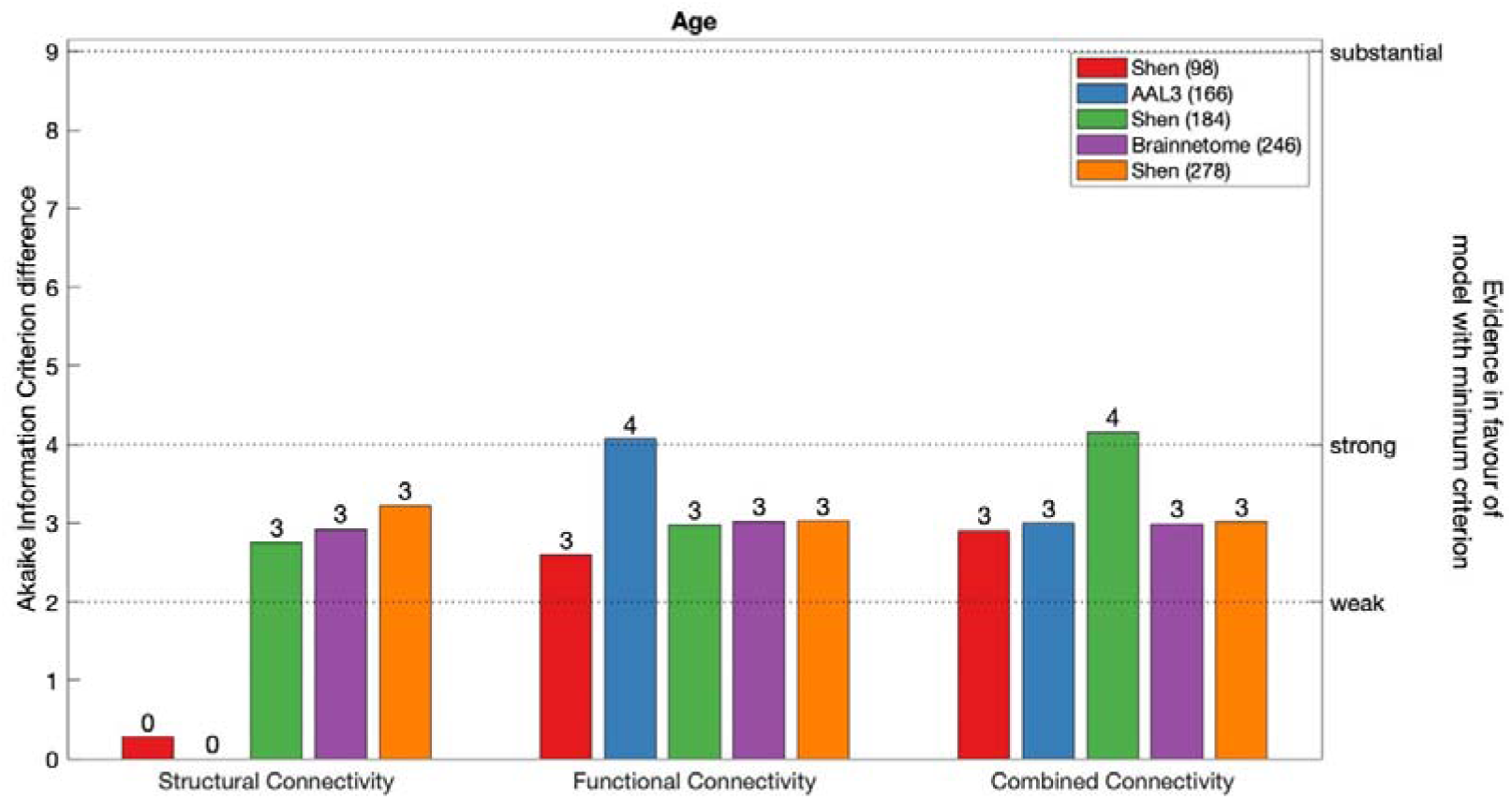
AIC difference for SC, FC and CC models of age, constructed with each parcellation scheme. Dotted line marks substantial evidence in favour of CC model defined with Shen (246) (minimum AIC across all modalities and parcellations).

**Figure 2.**
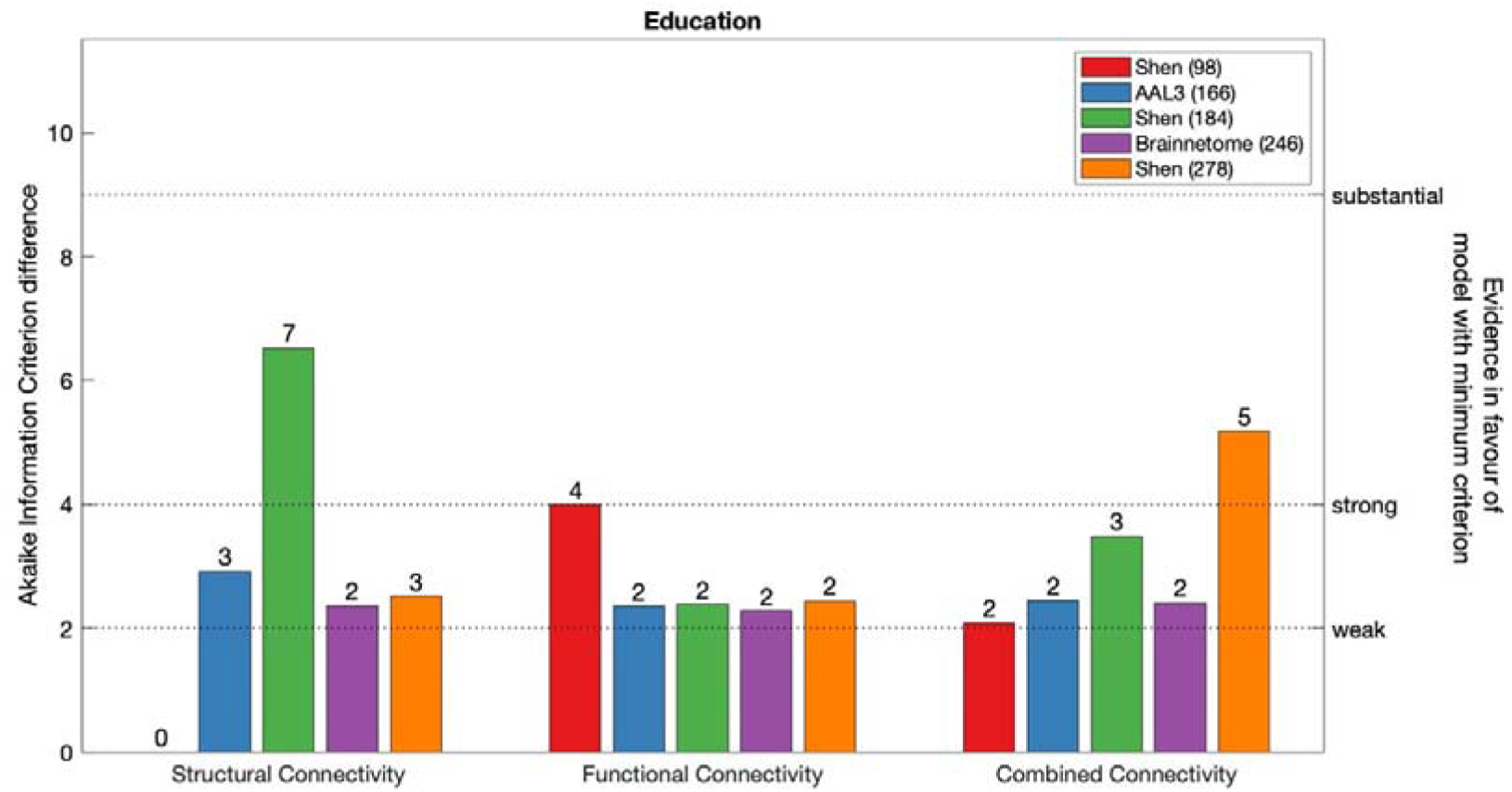
AIC difference for SC, FC and CC models of education, constructed with each parcellation scheme. Dotted line marks substantial evidence in favour of CC model defined with Shen (246) (minimum AIC across all modalities and parcellations).

**Figure 3.**
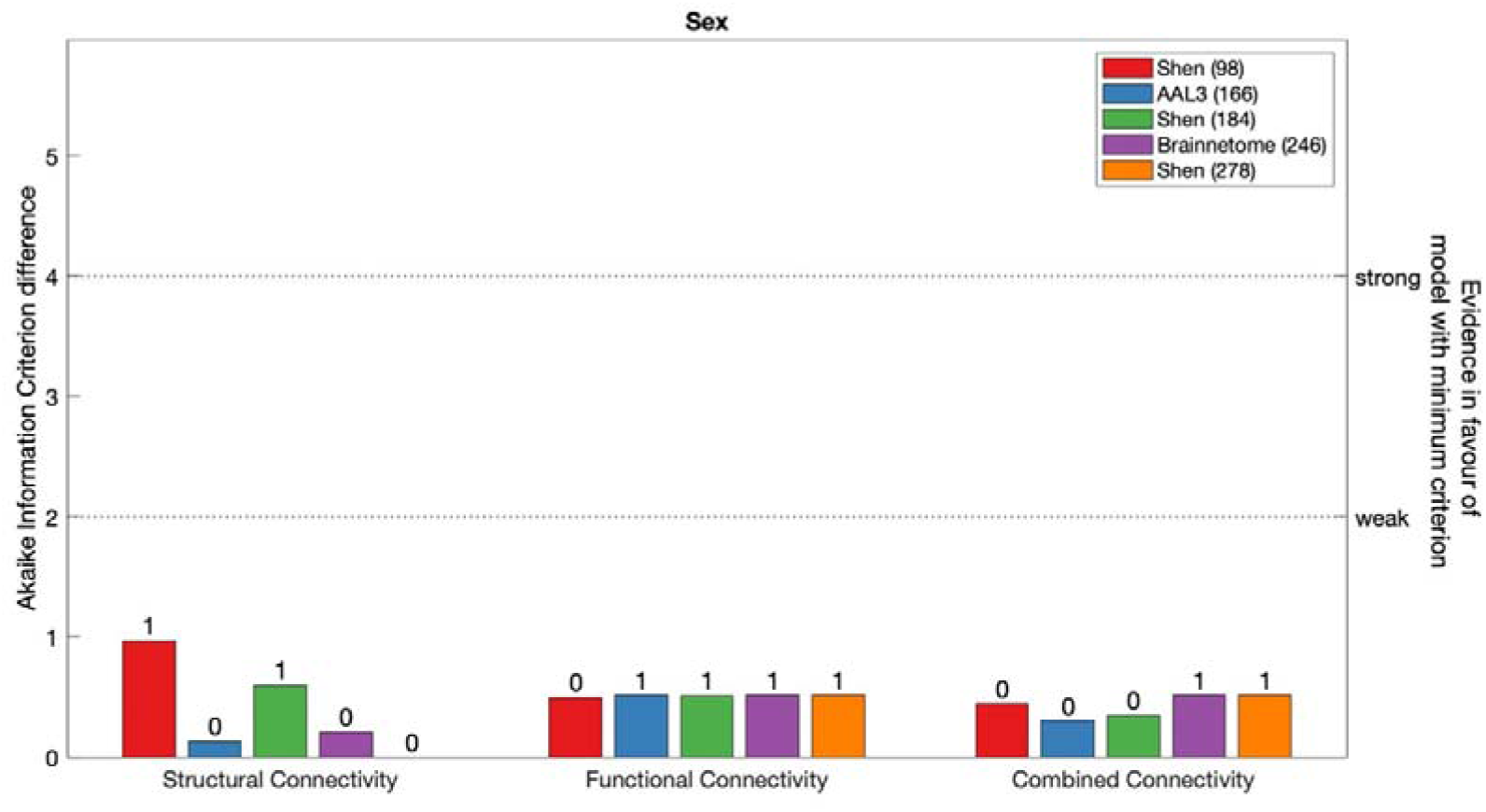
AIC difference for SC, FC and CC models of sex, constructed with each parcellation scheme. Dotted line marks substantial evidence in favour of CC model defined with Shen (246) (minimum AIC across all modalities and parcellations).

Next, across connectivity modalities and parcellations, education had lowest AIC value when modelled with SC defined with Shen (93). Within SC, there was evidence in favour of Shen (93) relative to AAL3 (166), Shen (278) and Brainnetome (246) and substantial evidence in favour of Shen (184). For FC, Brainnetome (246) was favoured to Shen (93). For CC, Shen (93) was favoured above Shen (184) and Shen (278).

Finally, there was no evidence to suggest that sex could be modelled more effectively using any single connectivity modality or parcellation.

##### Model comparison based on out-of-sample performance

Results of the cross-validation of models of demographics are presented in Figures 4-6. Cross-validation results demonstrated that only FC defined using Shen (278) parcellation did not produce above chance out-of-sample predictions of Education. All remaining models of all demographic variables performed above chance regardless of the parcellation schemes or the connectivity modality used as predictor.

**Figure 4.**
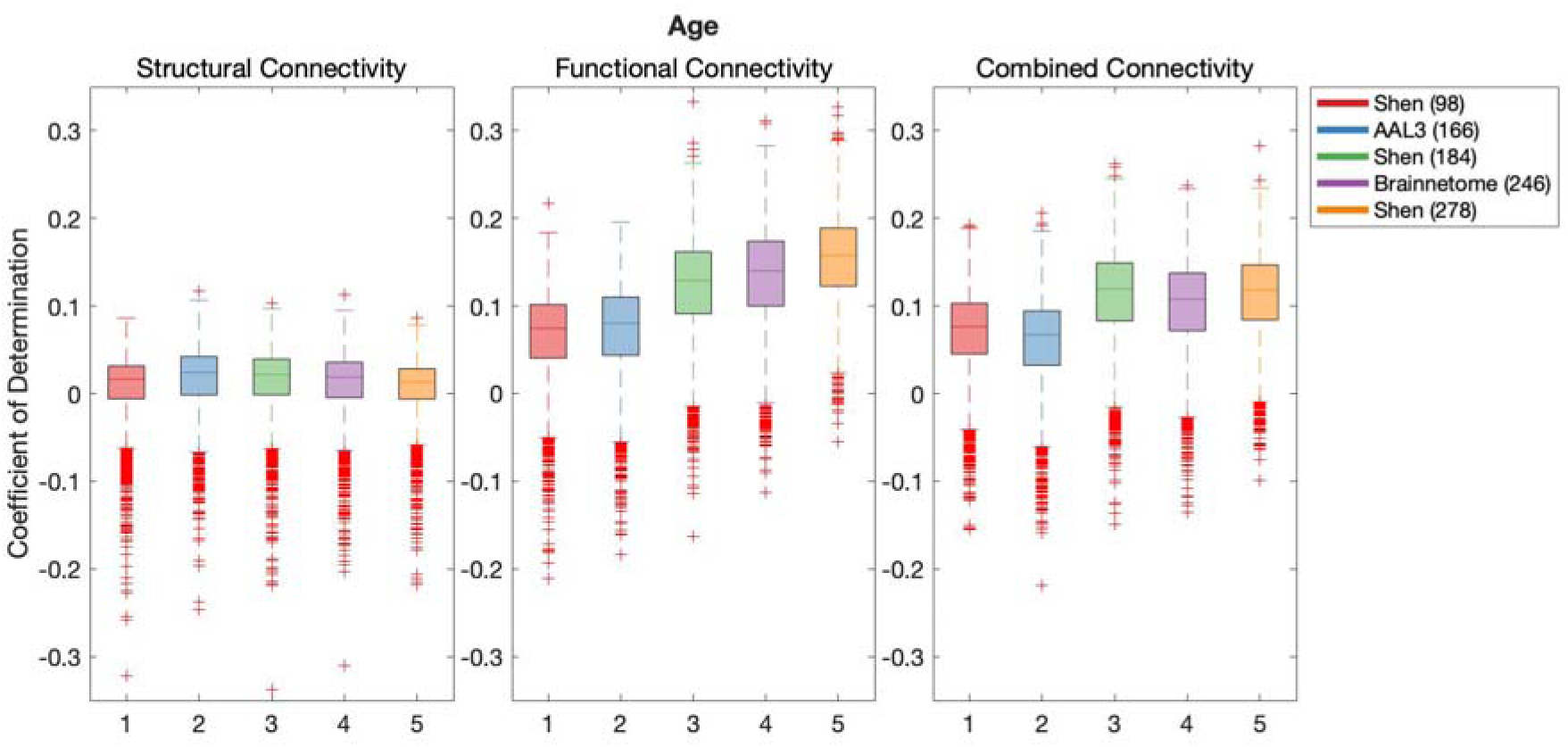
Results of BBC-CV of age models constructed with SC, FC and CC, as measured by the coefficient of determination. The solid lines show the median scores, the boxes show the interquartile range (IQR), and ticks outside of whiskers indicate outlier scores across all bootstrap samples. Filled boxes illustrate above-chance predictive performance and unfilled boxes illustrate bellow-chance prediction.

**Figure 5.**
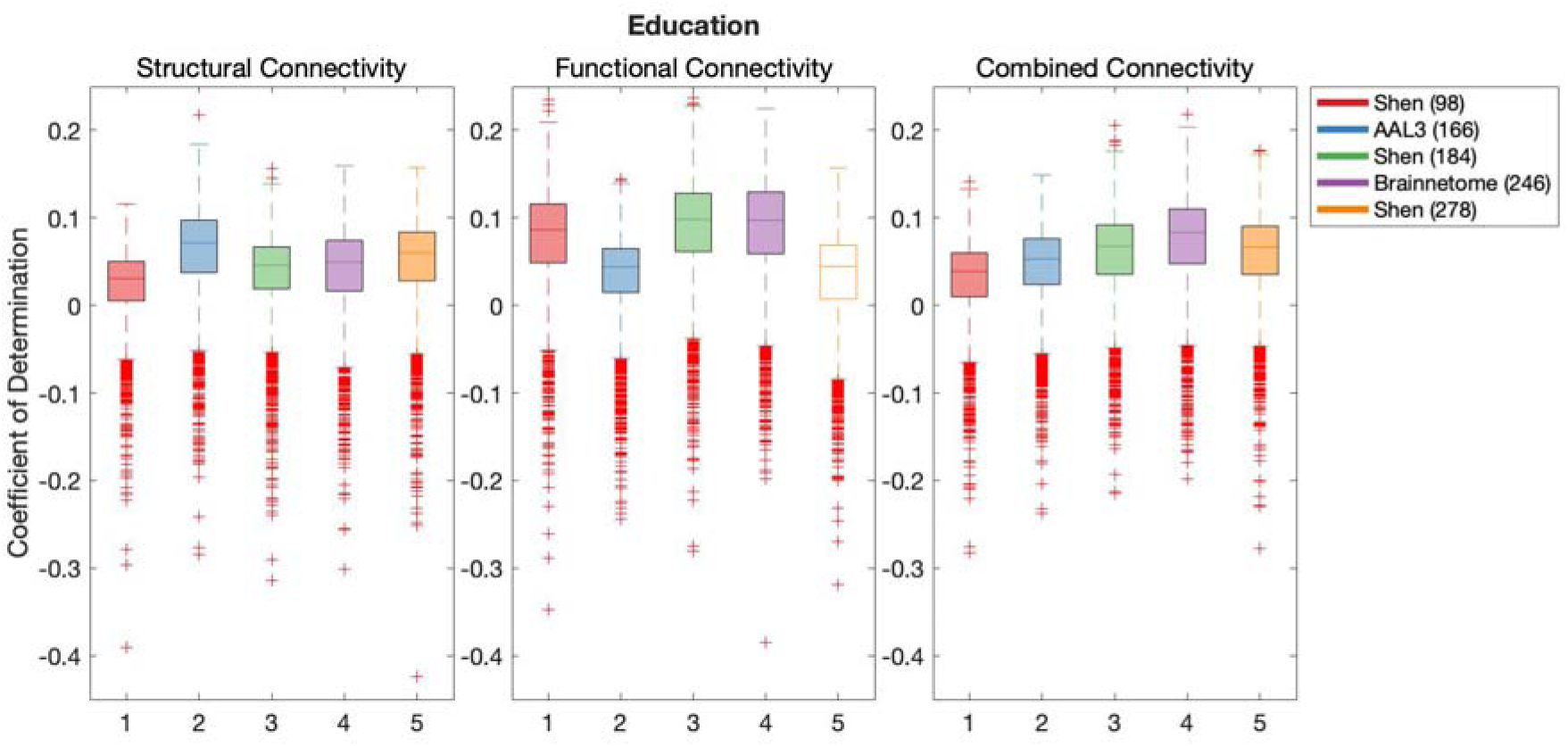
Results of BBC-CV of education models constructed with SC, FC and CC, as measured by the coefficient of determination. The solid lines show the median scores, the boxes show the interquartile range (IQR), and ticks outside of whiskers indicate outlier scores across all bootstrap samples. Filled boxes illustrate above-chance predictive performance and unfilled boxes illustrate bellow-chance prediction.

**Figure 6.**
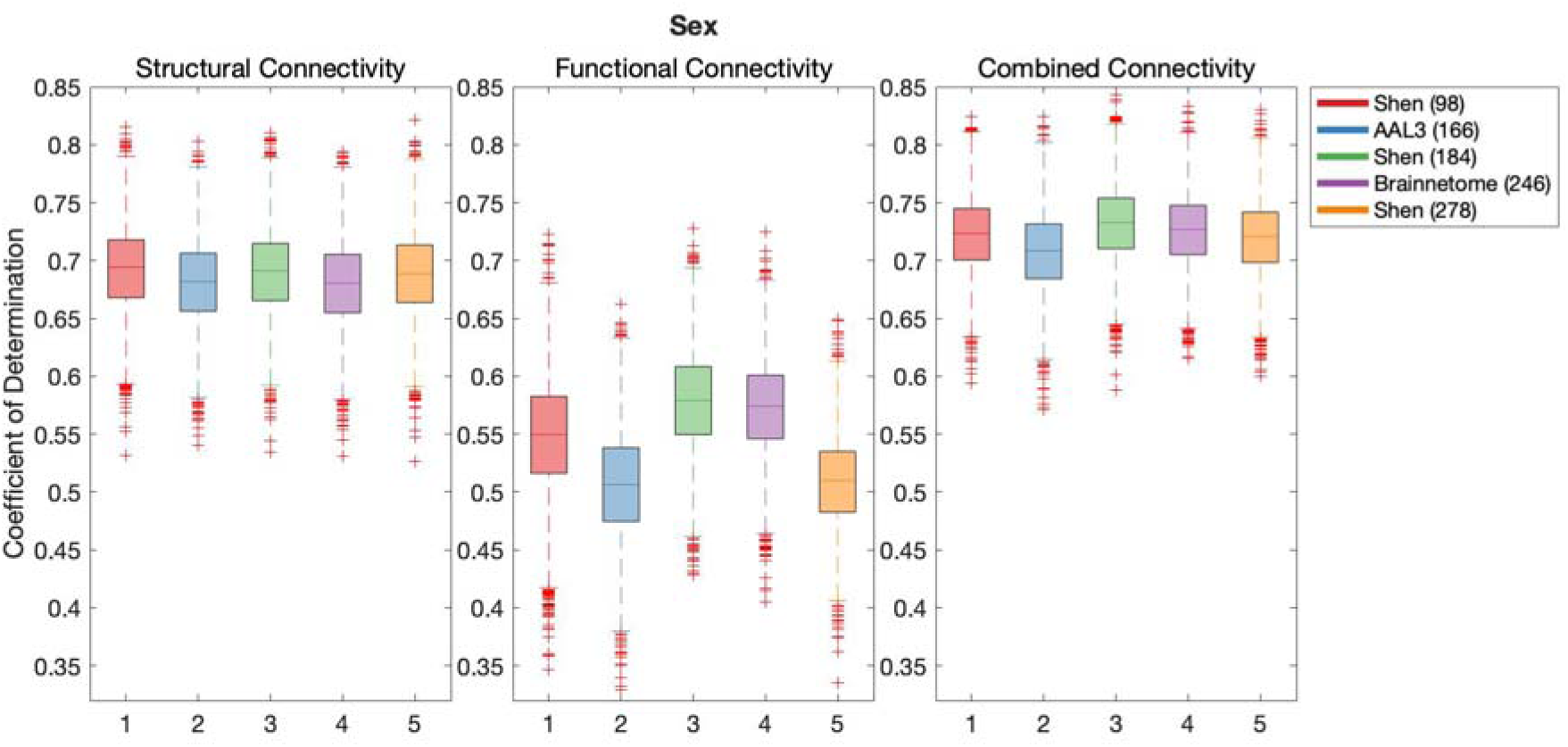
Results of BBC-CV of sex models constructed with SC, FC and CC, as measured by the coefficient of determination. The solid lines show the median scores, the boxes show the interquartile range (IQR), and ticks outside of whiskers indicate outlier scores across all bootstrap samples. Filled boxes illustrate above-chance predictive performance and unfilled boxes illustrate bellow-chance prediction.

Results of Wilcoxon rank sum test comparing out-of-sample models’ performance are shown in Table 1, where the False Discovery Rate (FDR) adjusted significance threshold equals 0.0095 (Benjamini & Hochberg, 1995)). The following results and effect sizes (Rosenthal et al., 1994) have been written for significant pair-wise comparisons presented in Table 1.

**Table 1.** Pairwise comparison Z-scores of coefficients of determination of demographics generated during BBC-CVT. Parcellations presented in rows were used as the reference, therefore positive values reflect that the parcellation presented in a given row performs better than the parcellation presented in a given column, whereas negative values indicate that the parcellation presented in a given row performs poorer than the parcellation presented in a given column. Comparisons were only performed for parcellations that produce better predictions than chance. All Z-values that are presented in bold meet the FDR-corrected p-value of 0.0095.

| **Connectivity** |  | Shen<br>(93) | AAL3<br>(166) | Shen<br>(184) | Brainnetome<br>(246) | Shen<br>(278) |
| --- | --- | --- | --- | --- | --- | --- |
| **Structural** | Shen (93) | - | **-14.07** | **-10.6** | **-4.97** | **4.66** |
|  | AAL3 (166) | **14.07** | - | **3.76** | **9.27** | **18.47** |
|  | Shen (184) | **10.6** | **-3.76** | - | **5.65** | **15.22** |
|  | Brainnetome<br>(246) | **4.97** | **-9.27** | **-5.65** | - | **9.61** |
|  | Shen (278) | **-4.66** | **-18.47** | **-15.22** | **-9.61** | - |
| **Functional** | Shen (93) | - | **-6.76** | **-50.22** | **-55.7** | **-70.37** |
|  | AAL3 (166) | **6.76** | - | **-44.4** | **-50.66** | **-65.95** |
|  | Shen (184) | **50.22** | **44.4** | - | **-10.03** | **-28.36** |
|  | Brainnetome<br>(246) | **55.7** | **50.66** | **10.03** | - | **-17.78** |
|  | Shen (278) | **70.37** | **65.95** | **28.36** | **17.78** | - |
| **Combined** | Shen (93) | - | **10.93** | **-41.68** | **-32.3** | **-42.57** |
|  | AAL3 (166) | **-10.93** | - | **-48.91** | **-40.62** | **-50.07** |
|  | Shen (184) | **41.68** | **48.91** | - | **11.47** | **0.62** |
|  | Brainnetome<br>(246) | **32.3** | **40.62** | **-11.47** | - | **-11.14** |
|  | Shen (278) | **42.57** | **50.07** | **-0.62** | **11.14** | - |

Education
| **Connectivity** |  | Shen<br>(93) | AAL3<br>(166) | Shen<br>(184) | Brainnetome<br>(246) | Shen<br>(278) |
| --- | --- | --- | --- | --- | --- | --- |
| **Structural** | Shen (93) | - | **-45.49** | **-21.13** | **-24.15** | **-35.74** |
|  | AAL3 (166) | **45.49** | - | **30.57** | **24.61** | **13.74** |
|  | Shen (184) | **21.13** | **-30.57** | - | **-5.43** | **-18.15** |
|  | Brainnetome<br>(246) | **24.15** | **-24.61** | **5.43** | - | **-11.95** |
|  | Shen (278) | **35.74** | **-13.74** | **18.15** | **11.95** | - |
| **Functional** | Shen (93) | - | **43.94** | **-11.91** | **-11.59** |  |
|  | AAL3 (166) | **-43.94** | - | **-52.78** | **-51.19** | - |
|  | Shen (184) | **11.91** | **52.78** | - | 0.02 | - |
|  | Brainnetome<br>(246) | **11.59** | **51.19** | -0.02 | - | - |
|  | Shen (278) | - | - | - | - | - |
| **Combined** | Shen (93) | - | **-19.49** | **-34.74** | **-46.51** | **-33.69** |
|  | AAL3 (166) | **19.49** | - | **-17.51** | **-32.87** | **-16.18** |
|  | Shen (184) | **34.74** | **17.51** | - | **-17.27** | 1.49 |
|  | Brainnetome<br>(246) | **46.51** | **32.87** | **17.27** | - | **18.7** |
|  | Shen (278) | **33.69** | **16.18** | -1.49 | **-18.7** | - |

|  |  | Sex |  |  |  |  |
| --- | --- | --- | --- | --- | --- | --- |
| Connectivity |  | Shen<br>(93) | AAL3<br>(166) | Shen<br>(184) | Brainnetome<br>(246) | Shen<br>(278) |
| **Structural** | Shen (93) | - | **15.09** | **3.45** | **16.85** | **5.98** |
|  | AAL3 (166) | **-15.09** | - | **-11.97** | 1.91 | **-9.36** |
|  | Shen (184) | **-3.45** | **11.97** | - | **13.8** | **2.6** |
|  | Brainnetome<br>(246) | **-16.85** | -1.91 | **-13.8** | - | **-11.2** |
|  | Shen (278) | **-5.98** | **9.36** | **-2.6** | **11.2** | - |
| **Functional** | Shen (93) | - | **40.03** | **-30.34** | **-25.6** | **41.02** |
|  | AAL3 (166) | **-40.03** | - | **-63.69** | **-61.65** | **-3.16** |
|  | Shen (184) | **30.34** | **63.69** | - | **6.52** | **66.43** |
|  | Brainnetome<br>(246) | **25.6** | **61.65** | **-6.52** | - | **64.62** |
|  | Shen (278) | **-41.02** | **3.16** | **-66.43** | **-64.62** | - |
| **Combined** | Shen (93) | - | **21.11** | **-14.08** | **-4.77** | **4.12** |
|  | AAL3 (166) | **-21.11** | - | **-33.98** | **-25.74** | **-17.36** |
|  | Shen (184) | **14.08** | **33.98** | - | **9.53** | **18.19** |
|  | Brainnetome<br>(246) | **4.77** | **25.74** | **-9.53** | - | **8.96** |
|  | Shen (278) | **-4.12** | **17.36** | **-18.19** | **-8.96** | - |

Out-of-sample predictive performance of age was significantly higher for models using AAL3 (166) parcellation in SC than any other parcellation, for Shen (278) in FC and for Shen (184) in CC. In the case of education, out-of-sample predictive performance was also significantly higher for models using AAL3 (166) parcellation in SC than alternative parcellations. In FC, highest out-of-sample predictive performance for education was found for both Shen (184) and Brainnetome (246), and there was no significant difference between their prediction performance. For CC, Brainnetome (246) parcellation performed more effectively at predicting education than alternative parcellations. Finally, Sex was most effectively predicted using Shen (93) parcellation in SC, and Shen (184) in both FC and CC.

#### Predictive modelling of cognition

##### AIC-based model comparison

Figures 7-11 demonstrate AIC for models of cognition composed with SC, FC and CC defined with the 5 parcellation schemes.

**Figure 7.**
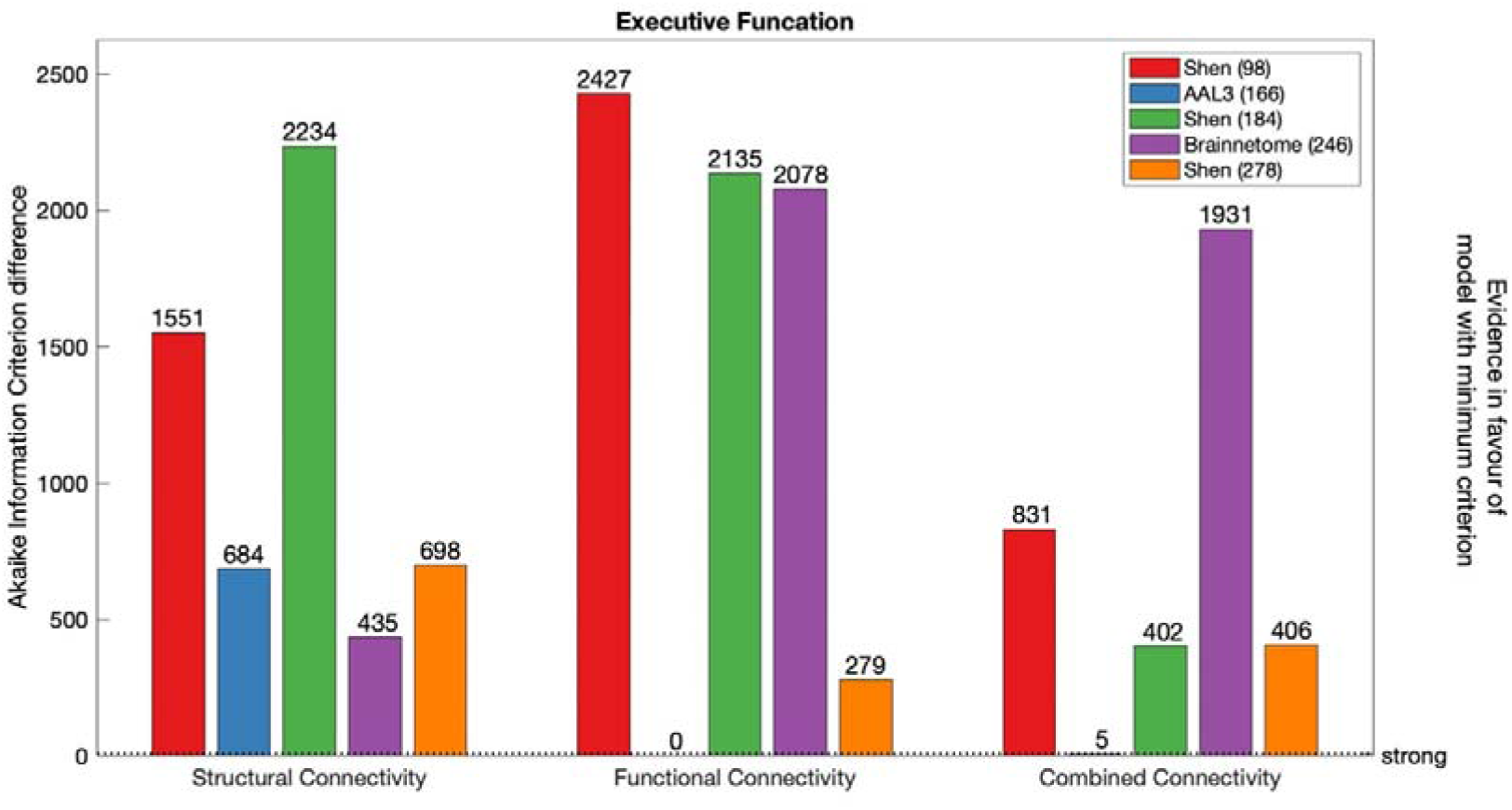
AIC difference for SC, FC and CC models of Executive Function, constructed with each parcellation scheme. Dotted line marks strong evidence in favour of FC model defined with AAL3 (166) (minimum AIC across all modalities and parcellations).

**Figure 8.**
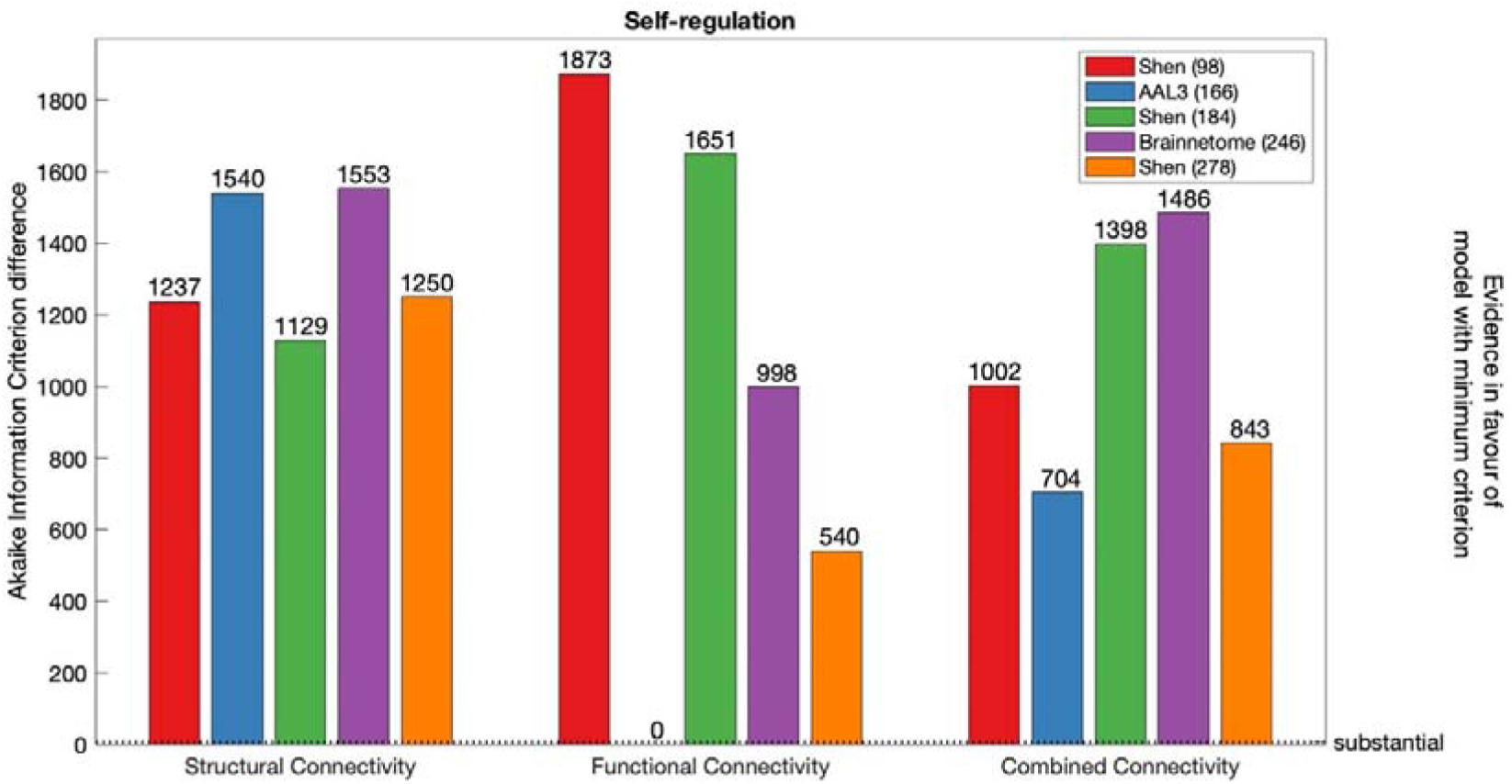
AIC difference for SC, FC and CC models of Self-regulation, constructed with each parcellation scheme. Dotted line marks substantial evidence in favour of FC model defined with AAL3 (166) (minimum AIC across all modalities and parcellations).

**Figure 9.**
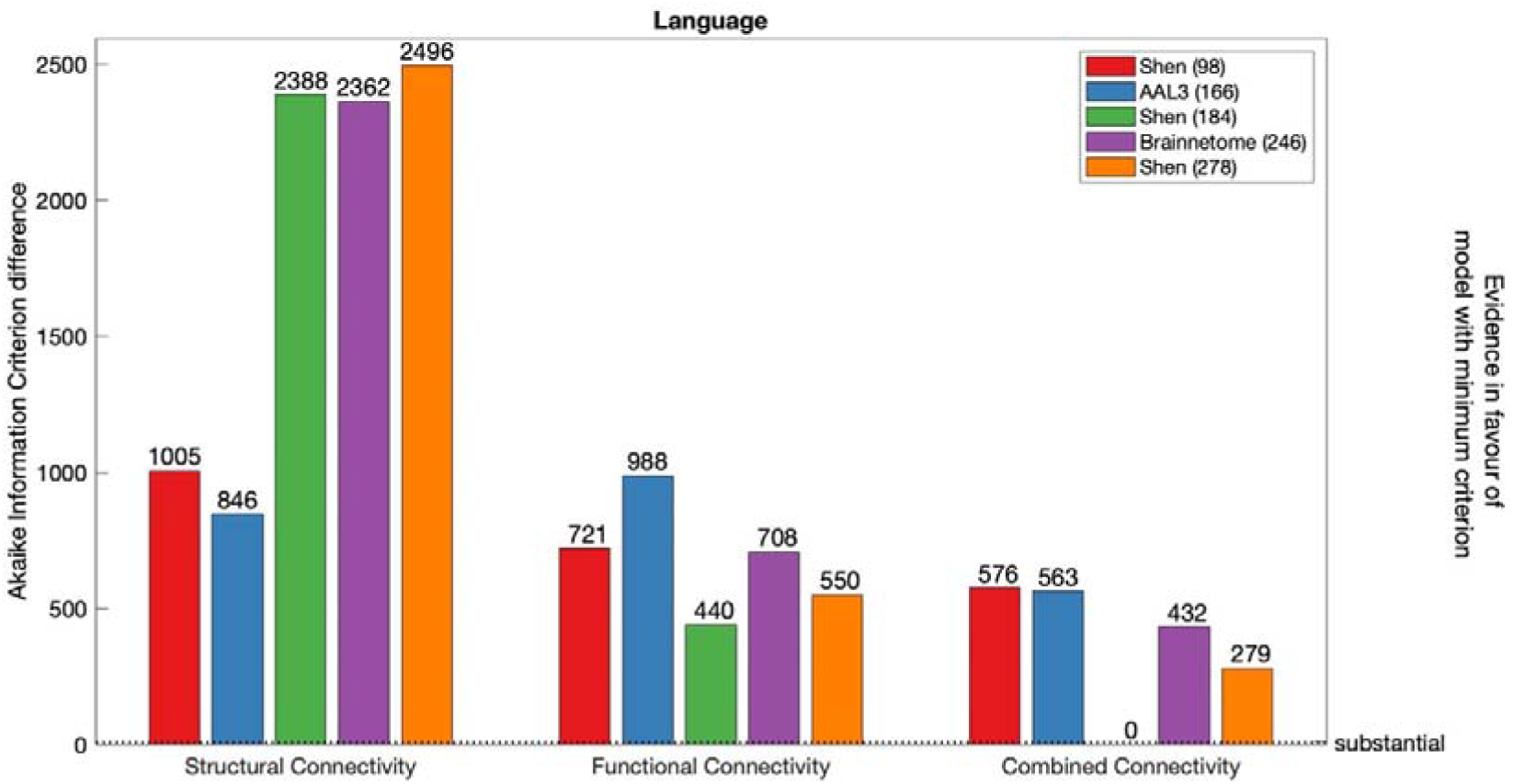
AIC difference for SC, FC and CC models of Language, constructed with each parcellation scheme. Dotted line marks substantial evidence in favour of CC model defined with Shen (246) (minimum AIC across all modalities and parcellations).

**Figure 10.**
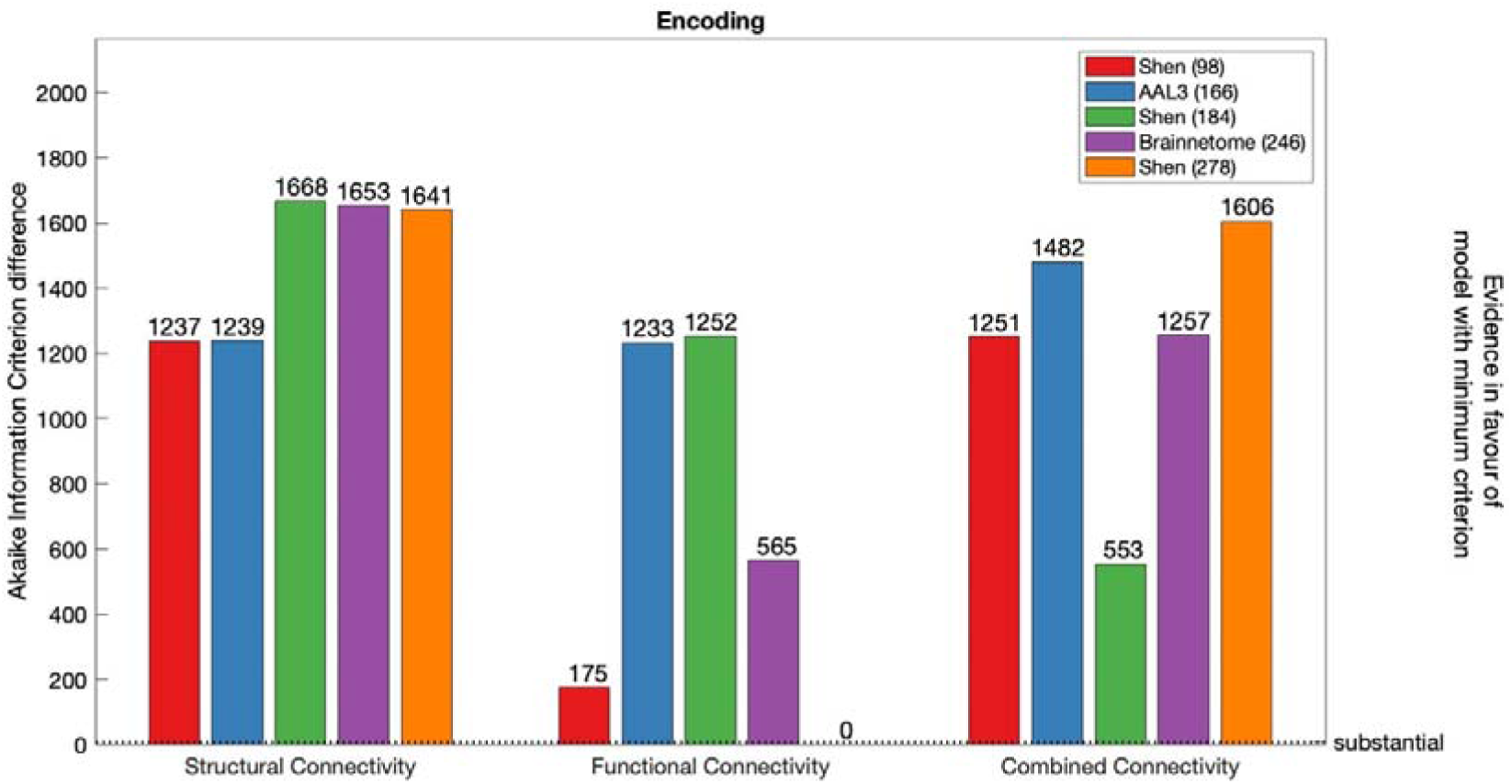
AIC difference for SC, FC and CC models of Encoding, constructed with each parcellation scheme. Dotted line marks substantial evidence in favour of FC model defined with Shen (278) (minimum AIC across all modalities and parcellations).

**Figure 11.**
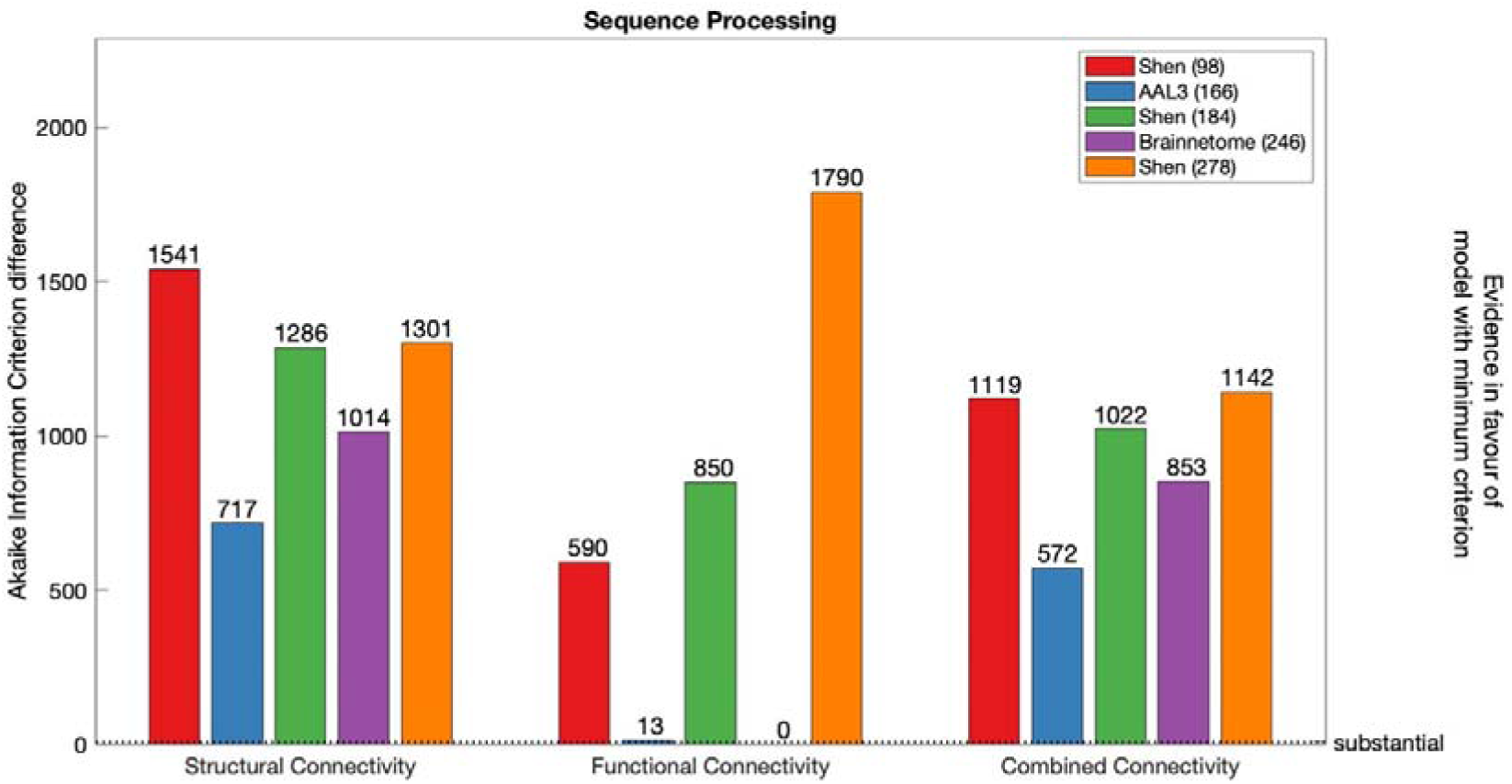
AIC difference for SC, FC and CC models of Encoding, constructed with each parcellation scheme. Dotted line marks substantial evidence in favour of FC model defined with Brainnetome (246) (minimum AIC across all modalities and parcellations).

AIC favoured models of Executive Function composed with FC defined with AAL3 (166) parcellation. For SC, there was substantial evidence in favour of Brainnetome (246) parcellation. For both FC and CC, there was substantial evidence in favour of AAL3 (166) parcellation.

There was substantial evidence in favour of FC model of Self-regulation constructed with AAL3 (166). In addition, for SC, there was substantial evidence in favour of Shen (184) parcellation. For CC, there was substantial evidence in favour of AAL3 (166) parcellation.

Overall, Language was best modelled by CC defined with Shen (184) parcellation. Similarly, when FC was used to model Language it was also best defined with Shen (184) parcellation. However, when FS was used to model Language it was best defined with AAL3 (166) parcellation.

Overall, Encoding was best modelled with FC defined with Shen (278) parcellation. When SC was used to model Encoding, best models were achieved with AAL3 (166) and Shen (93). When CC was used to model Encoding, according to AIC differences, the preferred model was defined with Shen (184).

Finally, across connectivity modalities and parcellations, Sequence Processing was best modelled with Brainnetome (246). When either SC or CC was used, AIC values were in favour of models obtained with AAL3 (166).

##### Model comparison based on out-of-sample performance

Results of the cross-validation of models of cognition are presented in Figures 12-17. It was found that the Shen (93) parcellation consistently produced out-of-sample predictions of cognition that explained more variance in unseen samples than chance. This was observed across SC, FC and CC.

**Figure 12.**
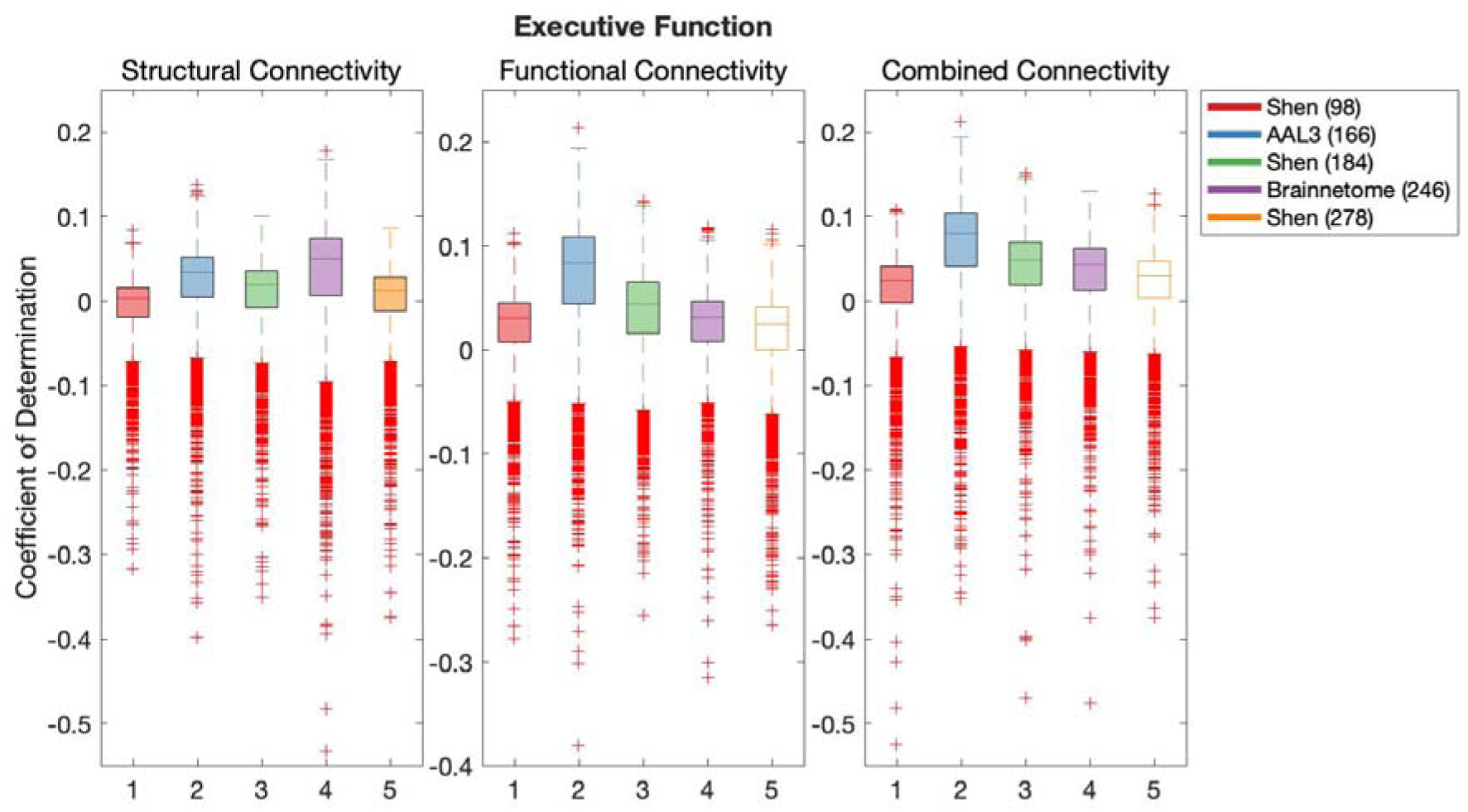
Results of BBC-CV of Executive Function models constructed with SC, FC and CC, as measured by the coefficient of determination. The solid lines show the median scores, the boxes show the interquartile range (IQR), and ticks outside of whiskers indicate outlier scores across all bootstrap samples. Filled boxes illustrate above-chance predictive performance and unfilled boxes illustrate bellow-chance prediction.

**Figure 13.**
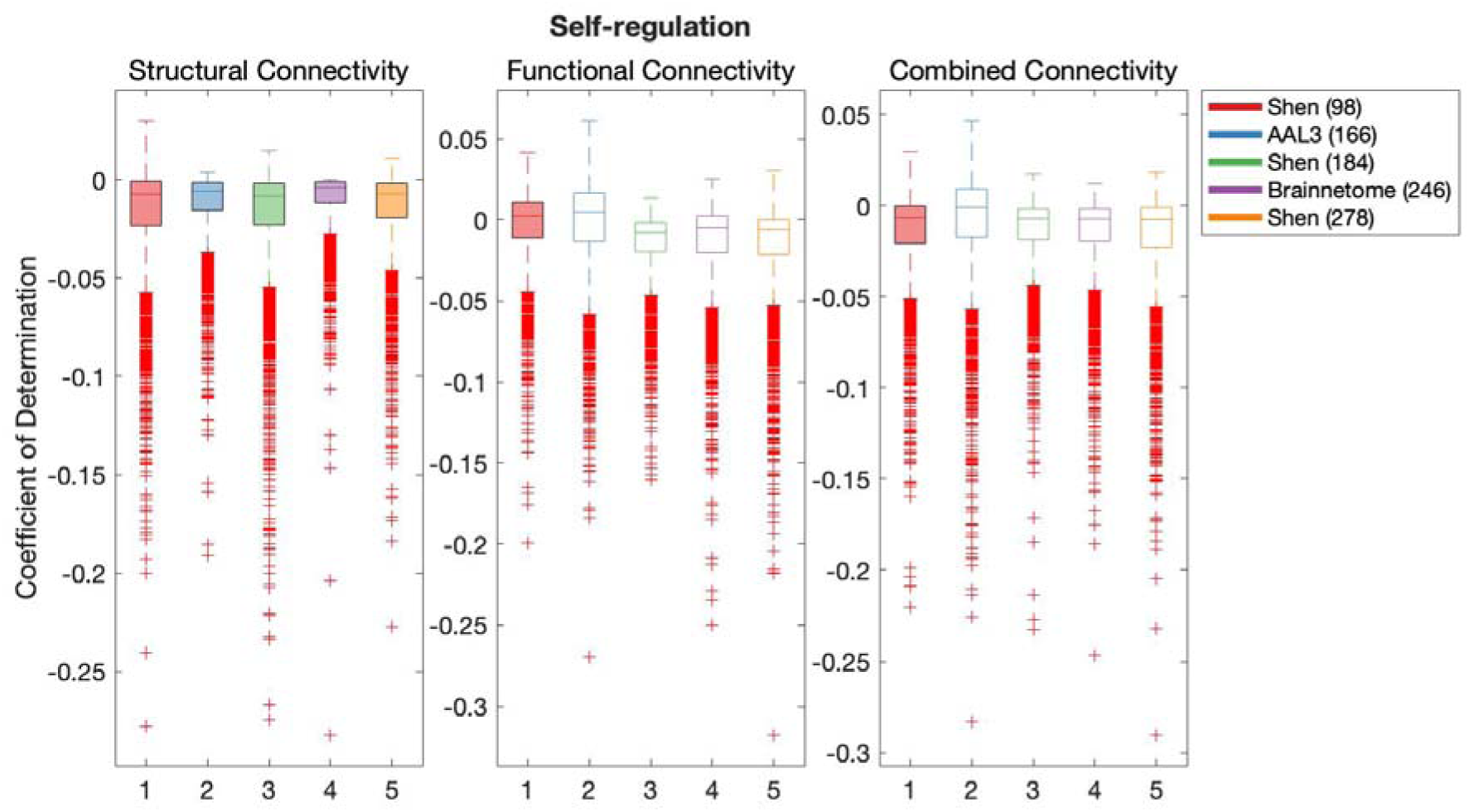
Results of BBC-CV of Self-regulation models constructed with SC, FC and CC, as measured by the coefficient of determination. The solid lines show the median scores, the boxes show the interquartile range (IQR), and ticks outside of whiskers indicate outlier scores across all bootstrap samples. Filled boxes illustrate above-chance predictive performance and unfilled boxes illustrate bellow-chance prediction.

**Figure 14.**
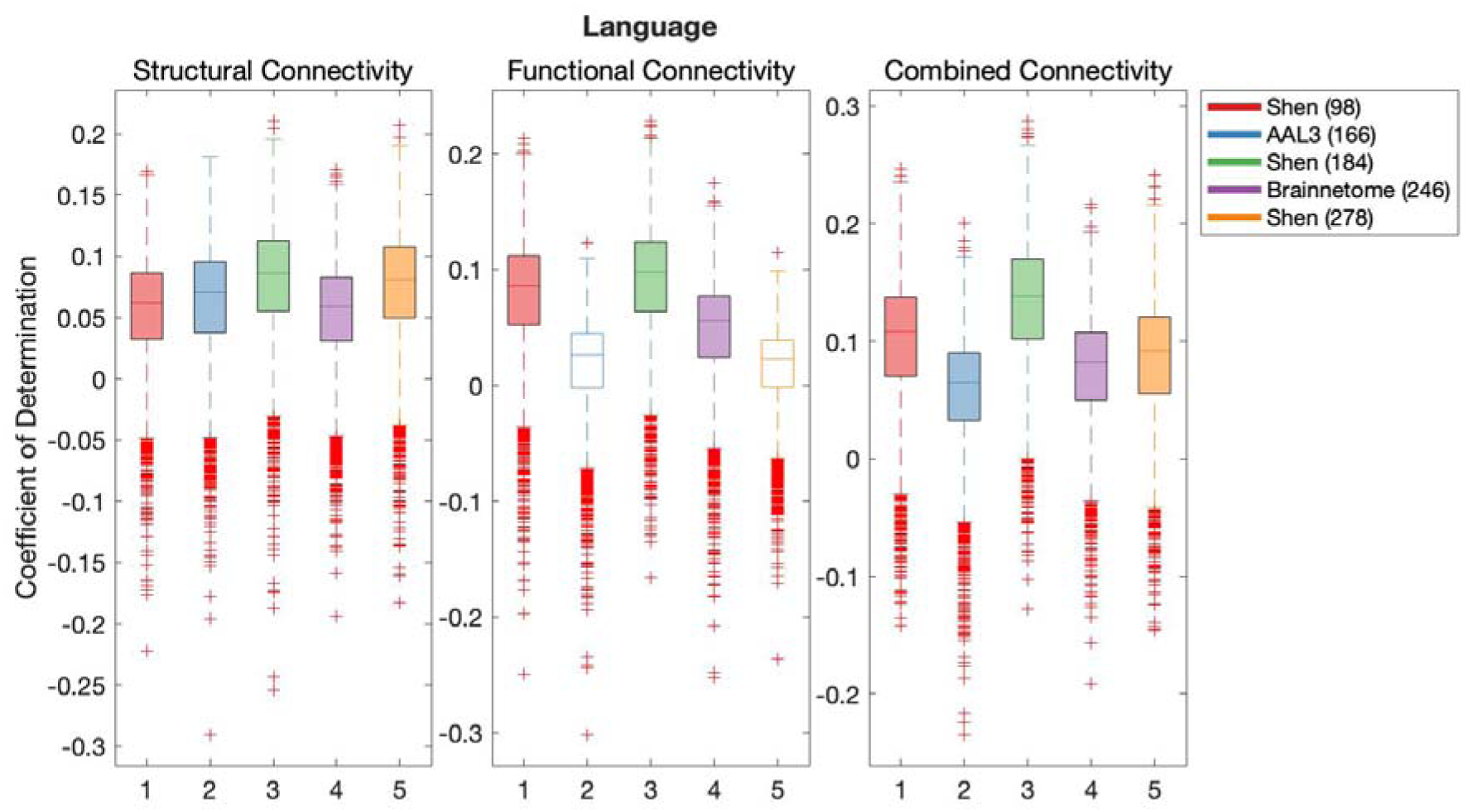
Results of BBC-CV of Language models constructed with SC, FC and CC, as measured by the coefficient of determination. The solid lines show the median scores, the boxes show the interquartile range (IQR), and ticks outside of whiskers indicate outlier scores across all bootstrap samples. Filled boxes illustrate above-chance predictive performance and unfilled boxes illustrate bellow-chance prediction.

**Figure 15.**
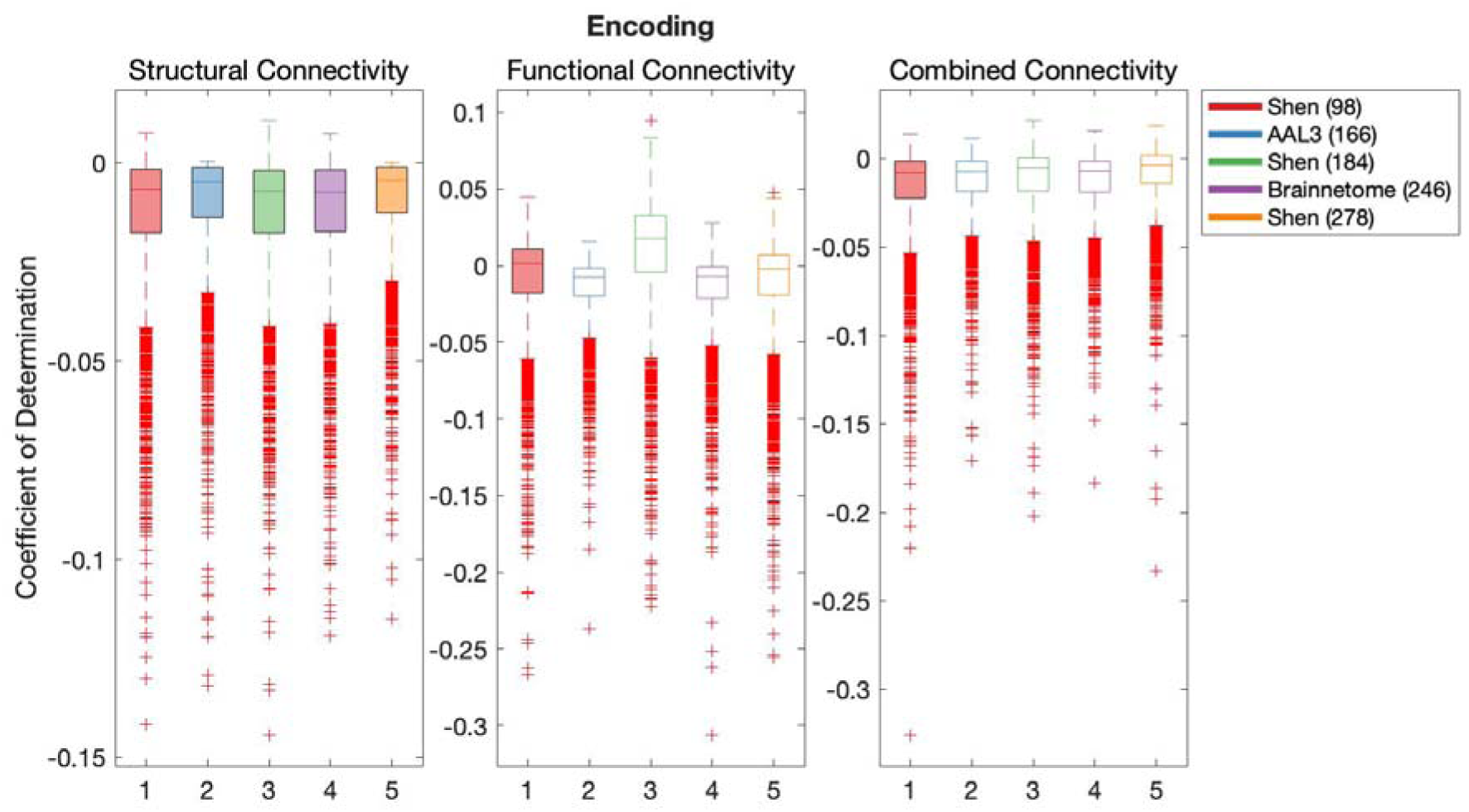
Results of BBC-CV of Encoding models constructed with SC, FC and CC, as measured by the coefficient of determination. The solid lines show the median scores, the boxes show the interquartile range (IQR), and ticks outside of whiskers indicate outlier scores across all bootstrap samples. Filled boxes illustrate above-chance predictive performance and unfilled boxes illustrate bellow-chance prediction.

**Figure 16.**
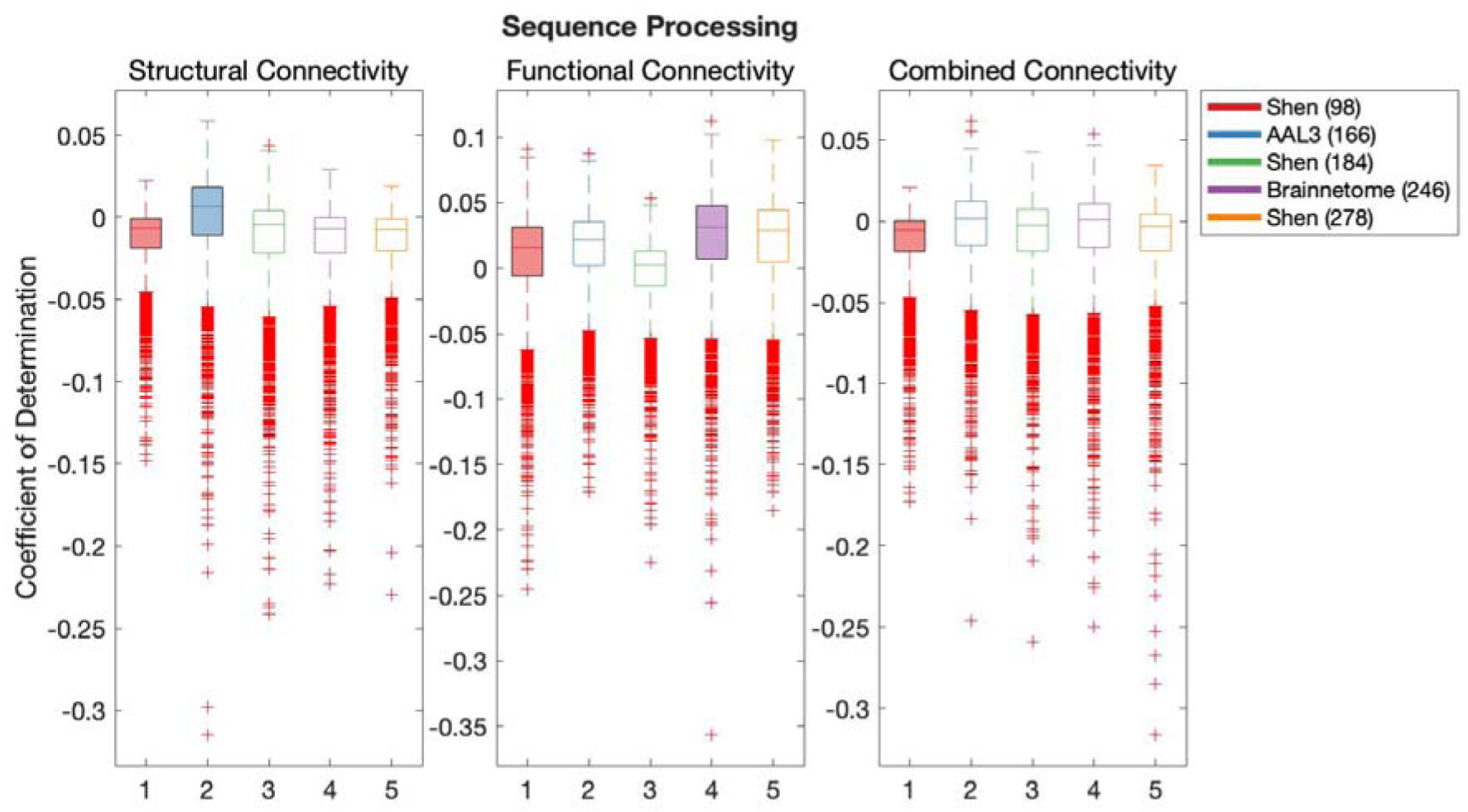
Results of BBC-CV of Sequence Processing models constructed with SC, FC and CC, as measured by the coefficient of determination. The solid lines show the median scores, the boxes show the interquartile range (IQR), and ticks outside of whiskers indicate outlier scores across all bootstrap samples. Filled boxes illustrate above-chance predictive performance and unfilled boxes illustrate bellow-chance prediction.

**Figure 17.**
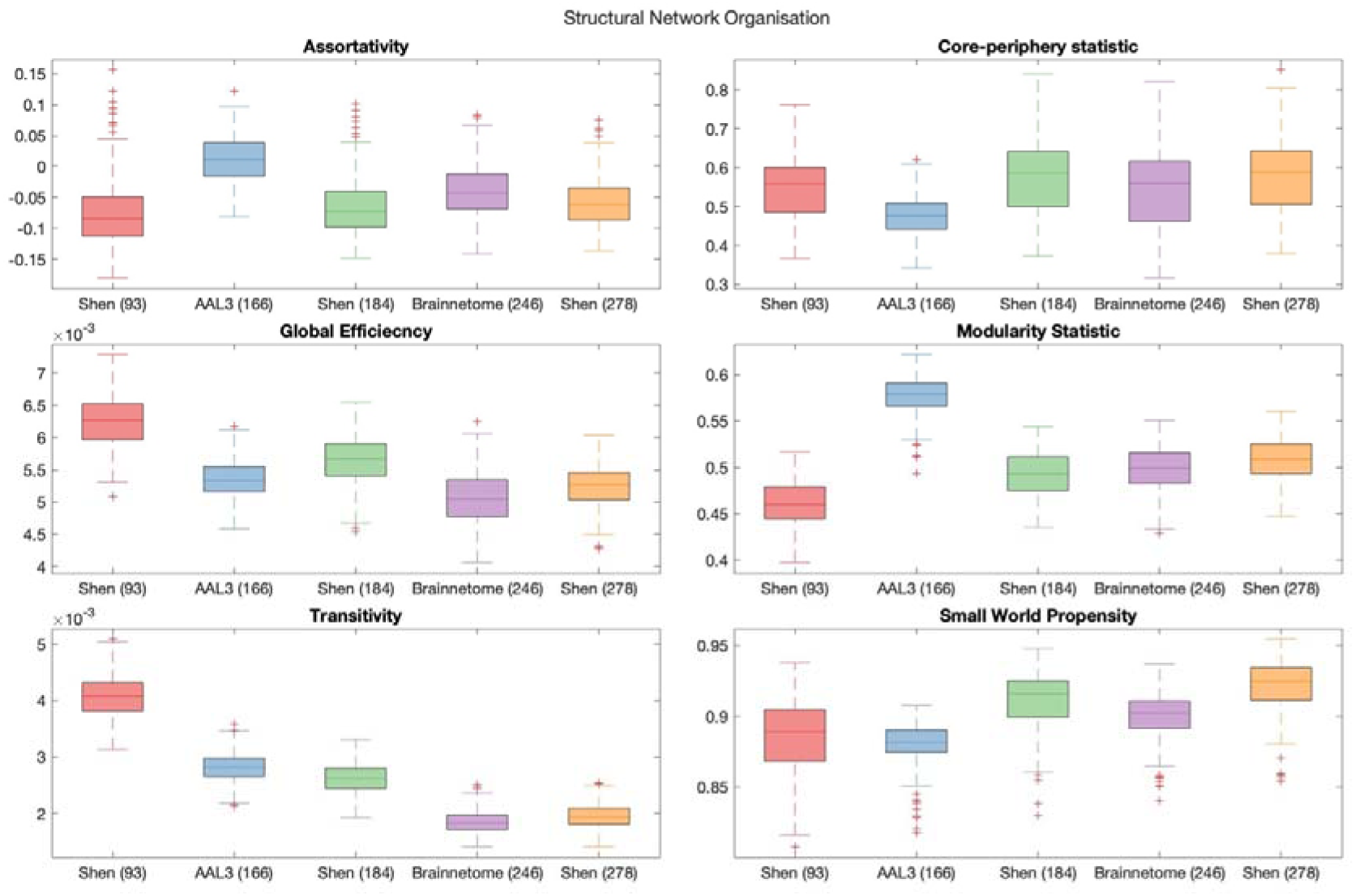
Graph theory measures of the global network organisation of SC.

All pairwise comparisons of the cross-validation performance are presented in Table 2. The probability of pairwise comparisons was adjusted with False Discovery Rate (Benjamini & Hochberg, 1995). All following comparisons have passed the significance threshold of 0.0184.

**Table 2.**
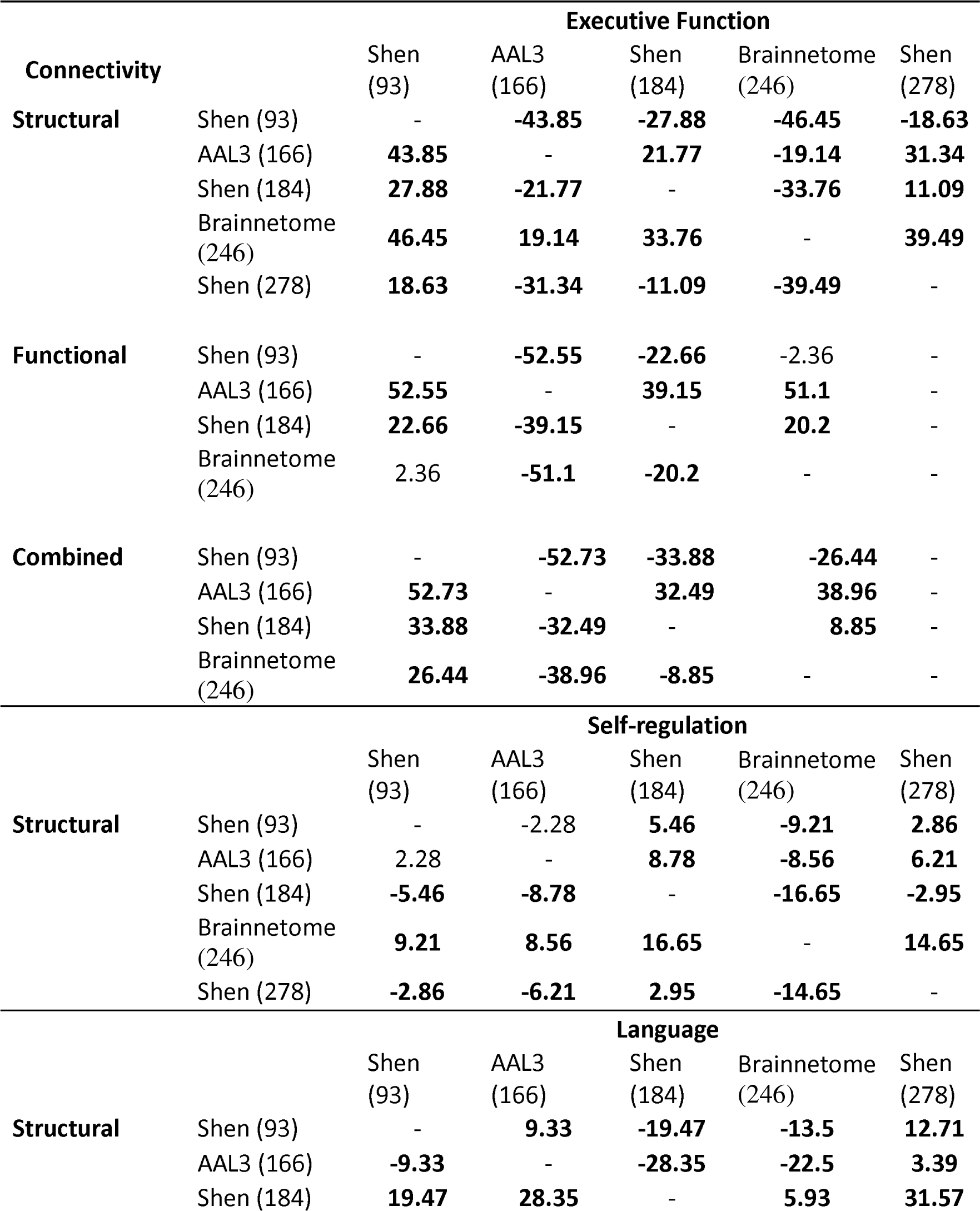
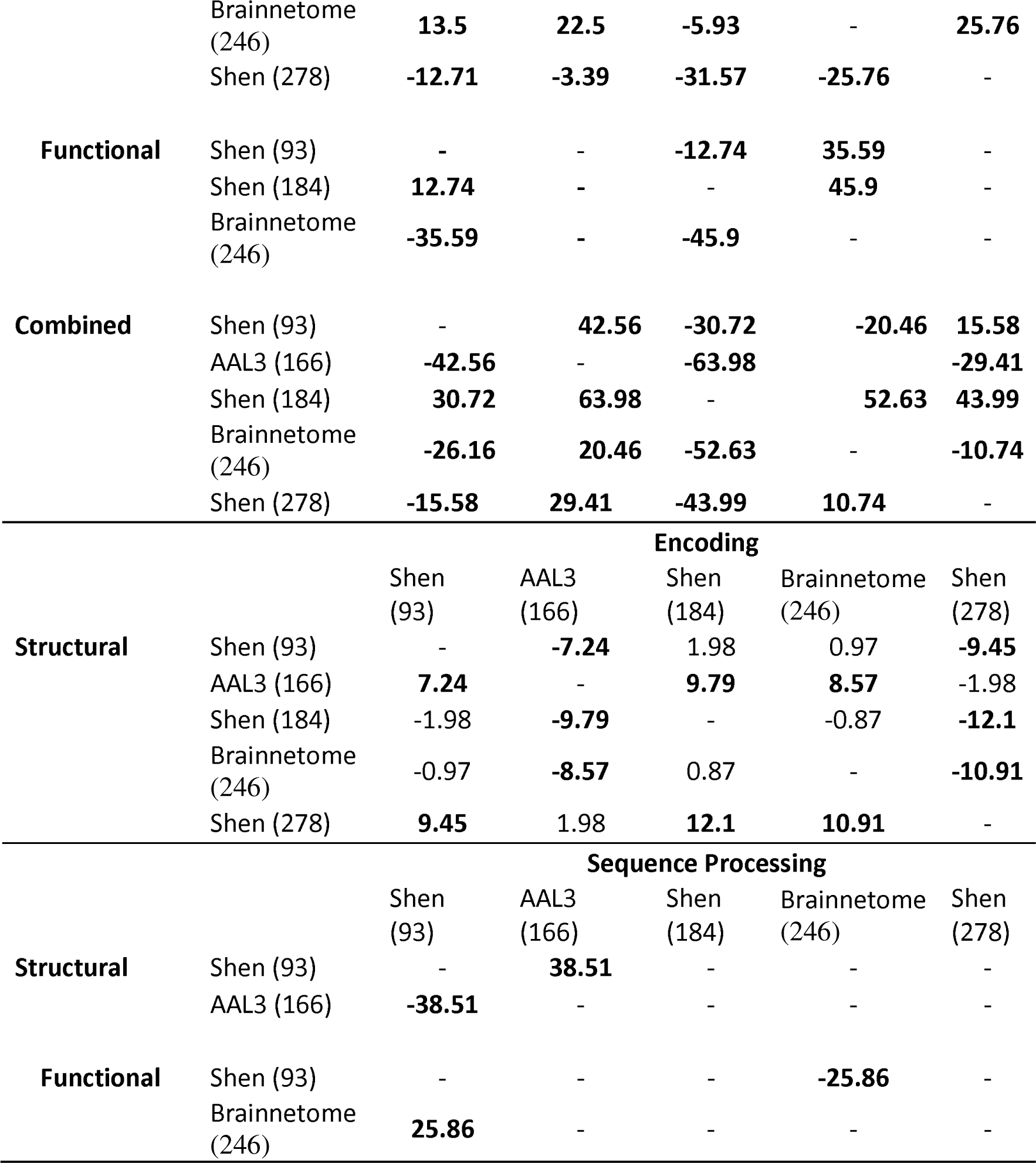
Pairwise comparison Z-scores of coefficients of determination of cognition generated during BBC-CVT. Parcellations presented in rows were used as the reference, therefore positive values reflect that the parcellation presented in a given row performs better than the parcellation presented in a given column, whereas negative values indicate that the parcellation presented in a given row performs poorer than the parcellation presented in a given column. Comparisons were only performed for parcellations that produce better predictions than chance. All Z-values that are presented in bold have the p-value <0.0184

Despite consistently explaining more variance in unseen sample than chance, Shen (93) rarely outperformed other parcellations at producing effective predictions of cognition (vs. Shen 184 Self-regulation structural connectivity, Language vs Brainnetome (246) structural connectivity, functional connectivity and combined connectivity, Language vs AAL3 (166) with CC and Language vs Shen 278 with CC). Next, AAL3 (166) consistently produced the most effective predictions of Executive Function utilising FC and CC. Similarly, Brainnetome (246) consistently produced the most effective predictions of Self-regulation utilising SC. Shen (184) parcellation consistently produced the most effective predictions of Language utilising FC. Shen (278) parcellation consistently produced the most effective predictions of Encoding utilising SC. There was no other parcellation scheme that consistently outperformed other parcellation schemes for any given type of connectivity or any cognitive construct.

### Graph theory

#### Comparison of global organisation across parcellations

Figures 17 and 18 show the distribution of scores obtained for each participant’s global graph theory measures for each parcellation. The following Tables 3 and 4 present pair-wise comparisons made to assess differences in global graph theory measures across parcellations. It was generally found that parcellation schemes significantly impacted network organisation, as measured by all 6 graph theory measures. However, the direction of this impact was not necessarily shared across SC and FC. For example, structural parcellation scheme (AAL3, 166) produced high modularity statistic in SC, but low modularity statistic in FC. Further, high resolution scheme (Shen, 278) produced lowest global efficiency and transitivity in SC, whereas in FC a lower resolution parcellation (AAL3, 166) produced lowest global efficiency and transitivity.

**Figure 18.**
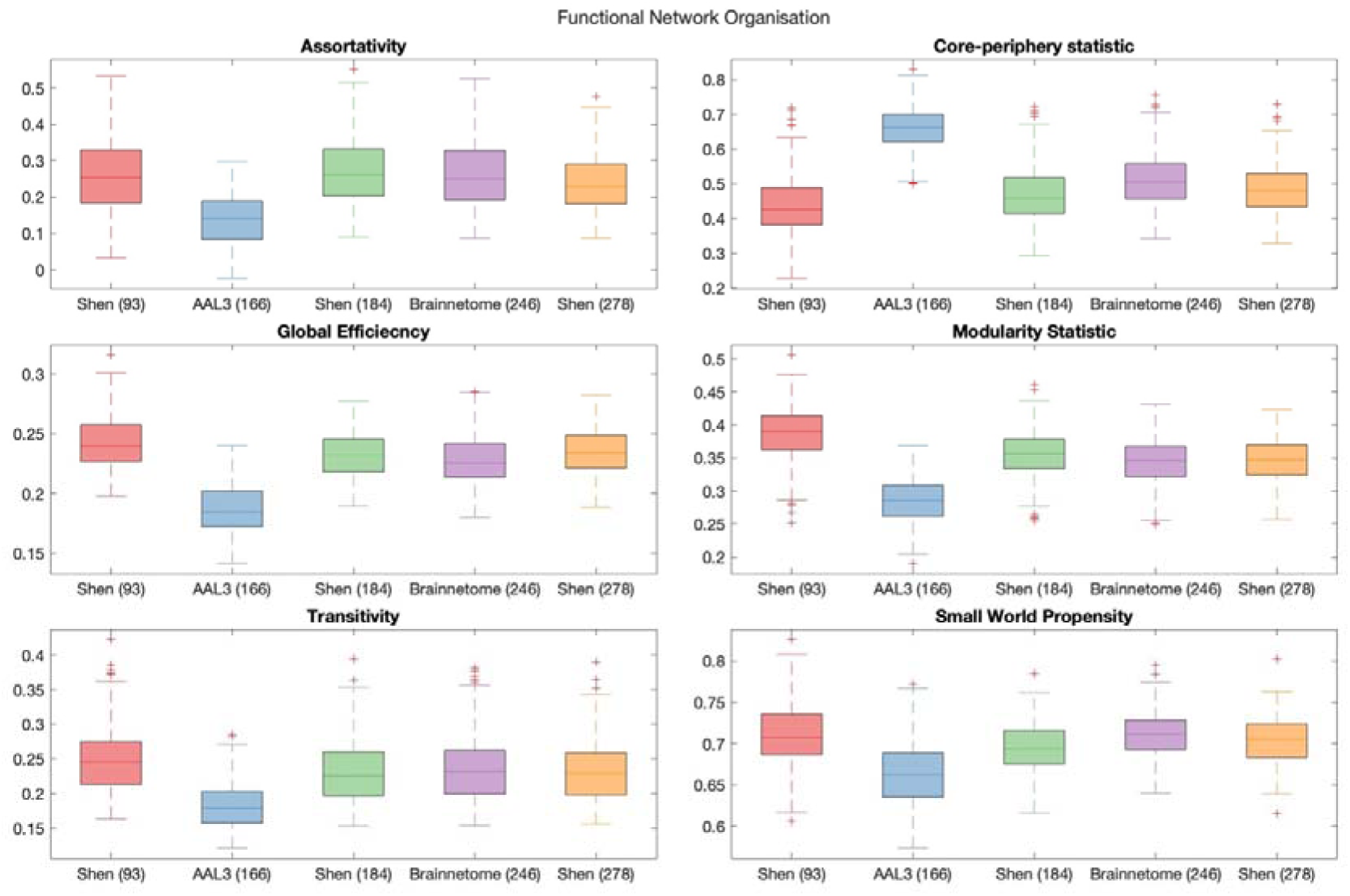
Graph theory measures of the global network organisation of FC.

**Table 3.**
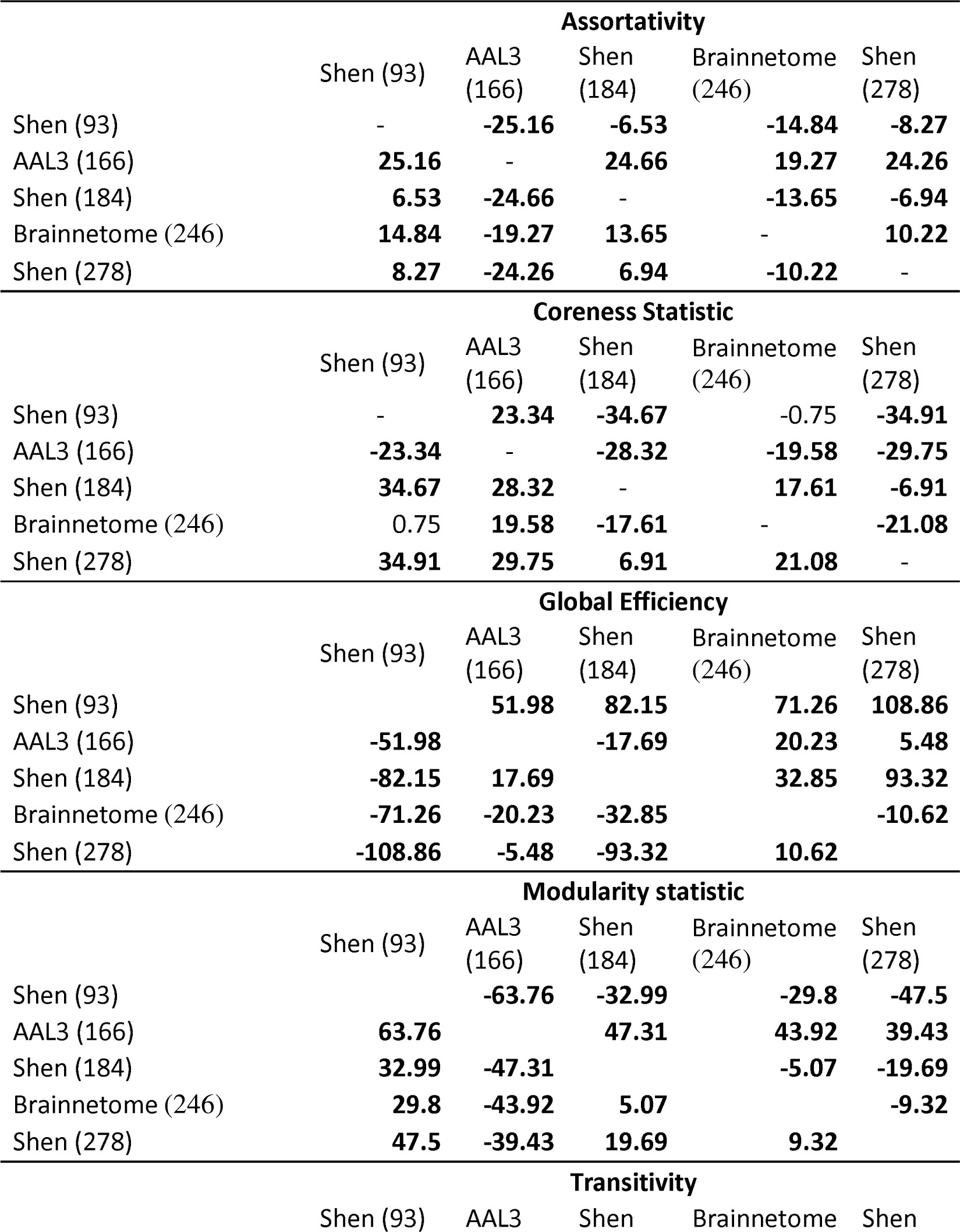
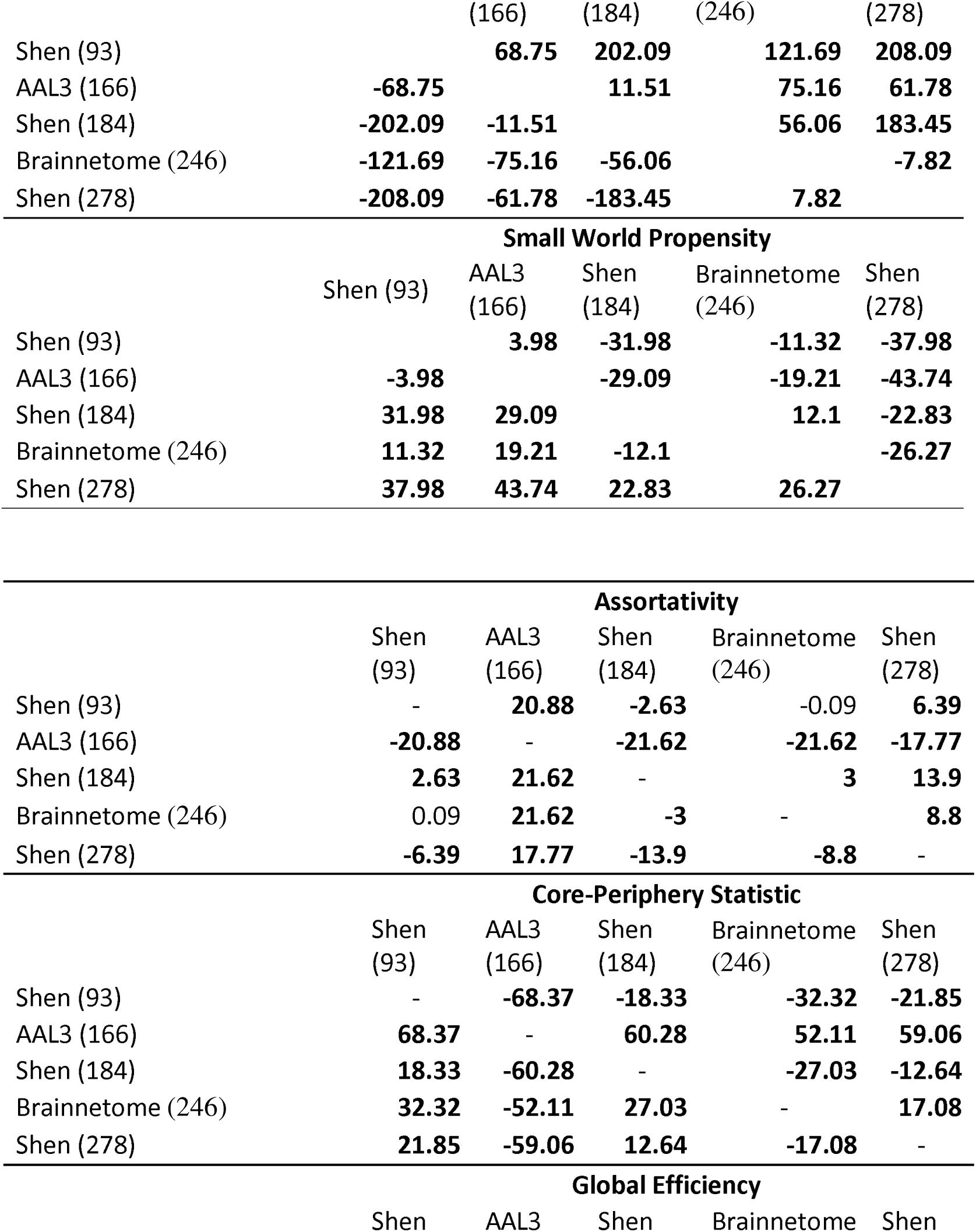
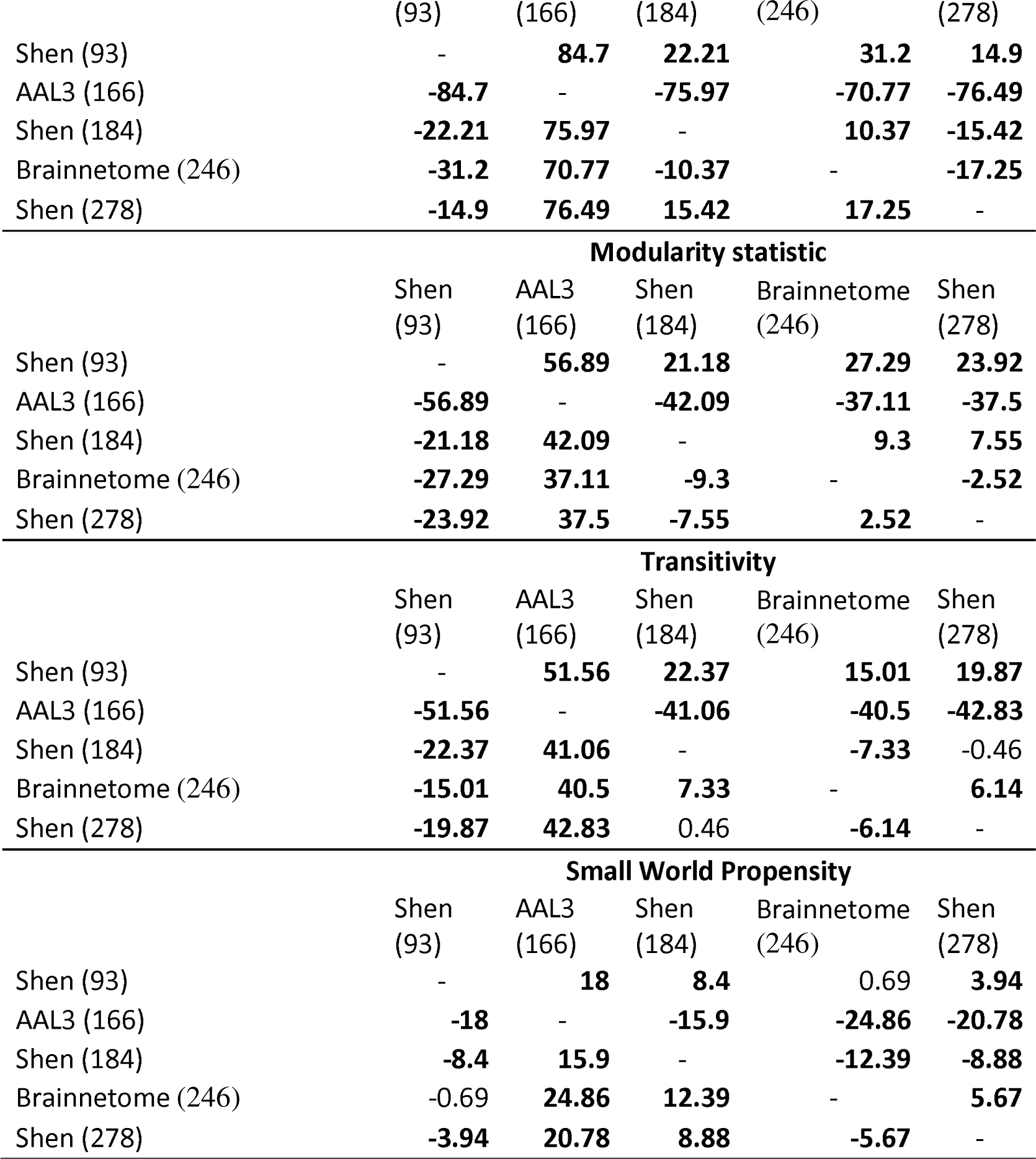
Pairwise comparison t-statistics of structural connectivity organisation yielded by each network parcellation scheme. Parcellations presented in rows were used as the reference, therefore positive values reflect that the parcellation presented in a given row has higher global property (assortativity, core-periphery statistic, global efficiency, modularity, transitivity, small world propensity) than the parcellation presented in a given column, whereas negative values indicate that the parcellation presented in a given row has lower global property than the parcellation presented in a given column. All t-scores presented in bold have passed the FDR-adjusted critical p-value =<.0275.

#### Predictive modelling of demographics

##### AIC-based model comparison

Figures 19-21 demonstrate AIC for models of demographics composed with global graph theory measures of SC, FC and CC when defined with the 5 parcellation schemes.

**Figure 19.**
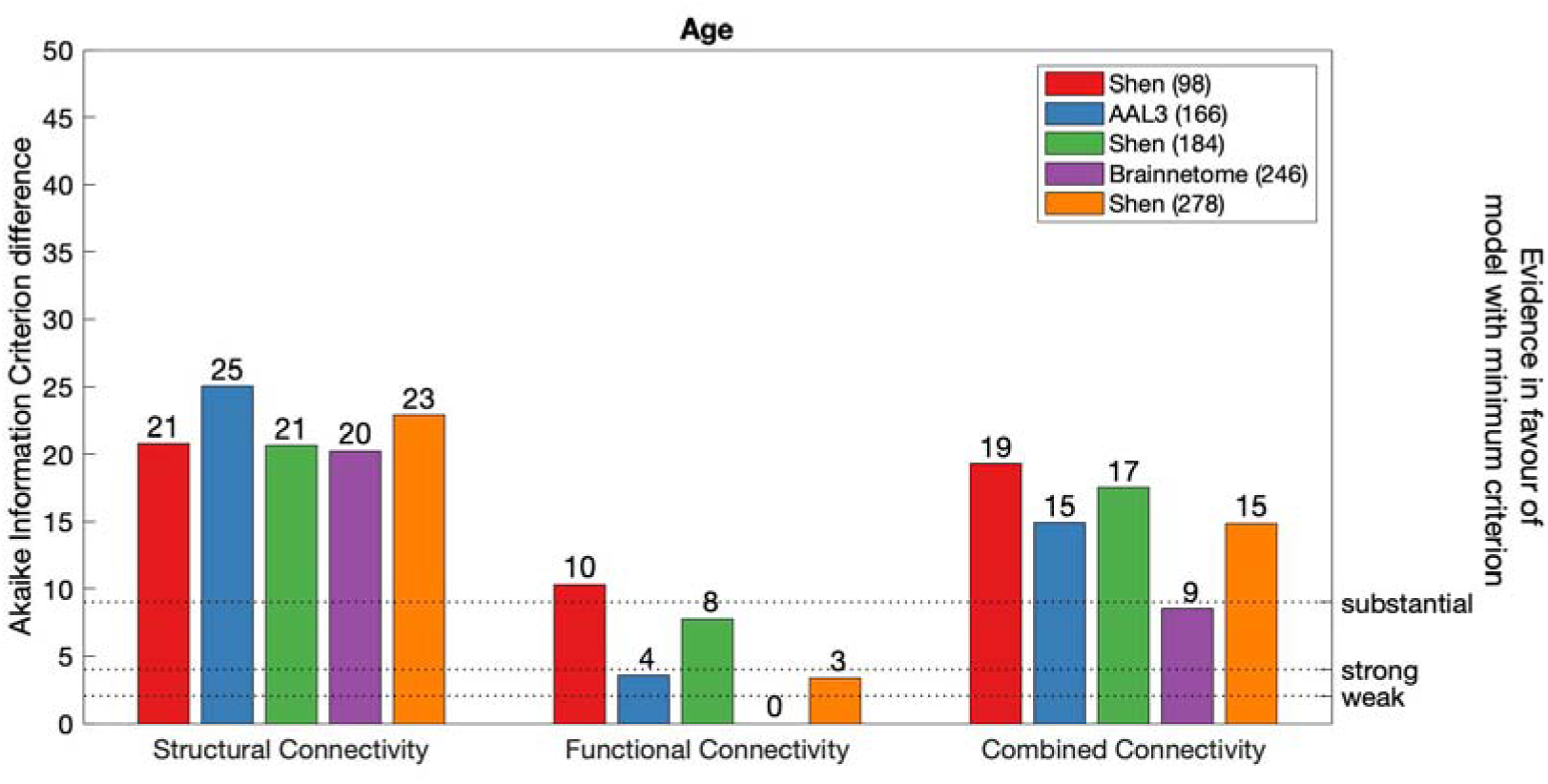
AIC difference for global graph theory measure models of Age, constructed with each parcellation scheme. Dotted line marks weak, strong and substantial evidence in favour of FC model defined with Brainnetome (246) (minimum AIC across all modalities and parcellations).

**Figure 20.**
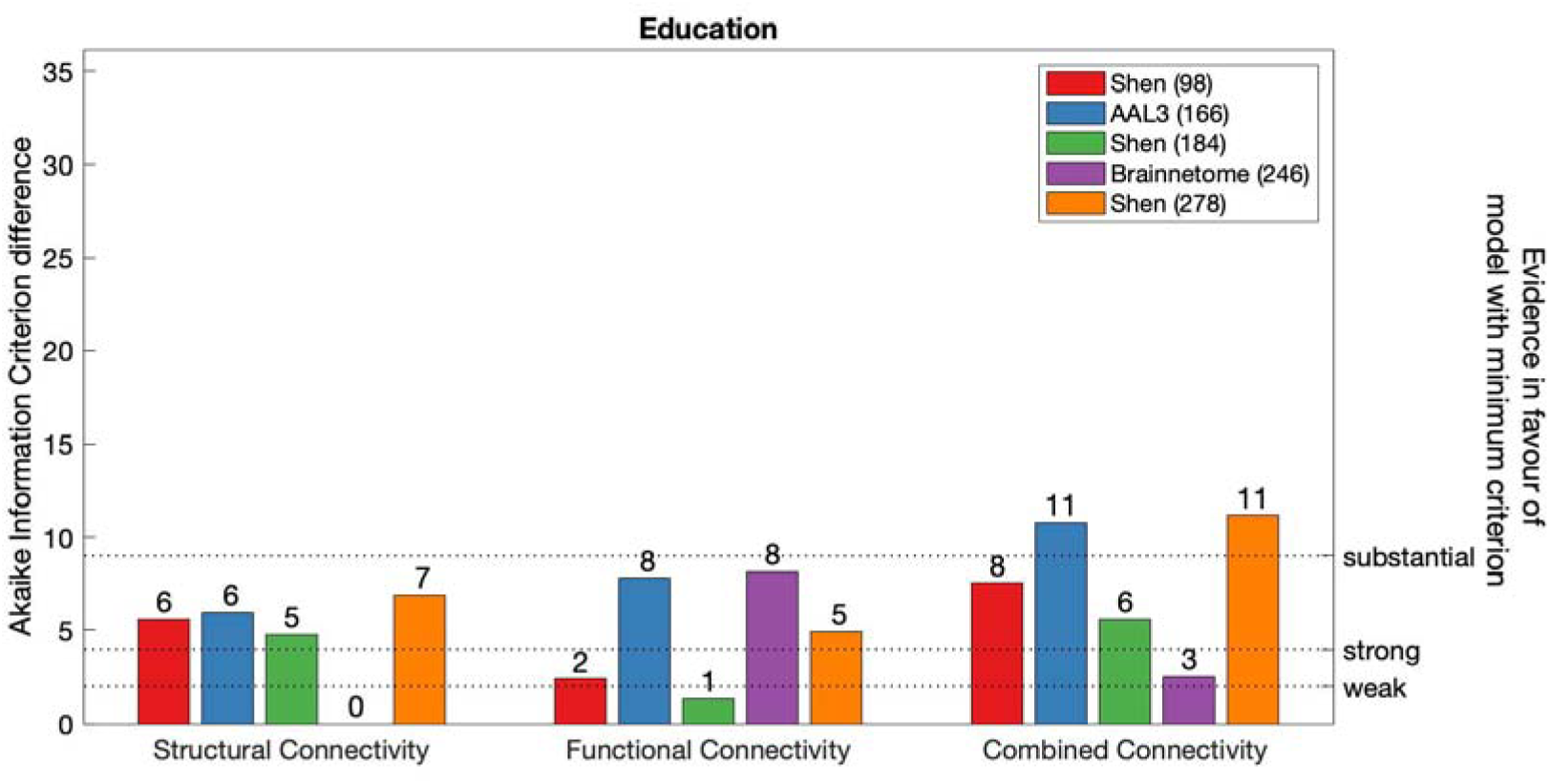
AIC difference for global graph theory measure models of Education, constructed with each parcellation scheme. Dotted line marks weak, strong and substantial evidence in favour of SC model defined with Brainnetome (246) (minimum AIC across all modalities and parcellations).

**Figure 21.**
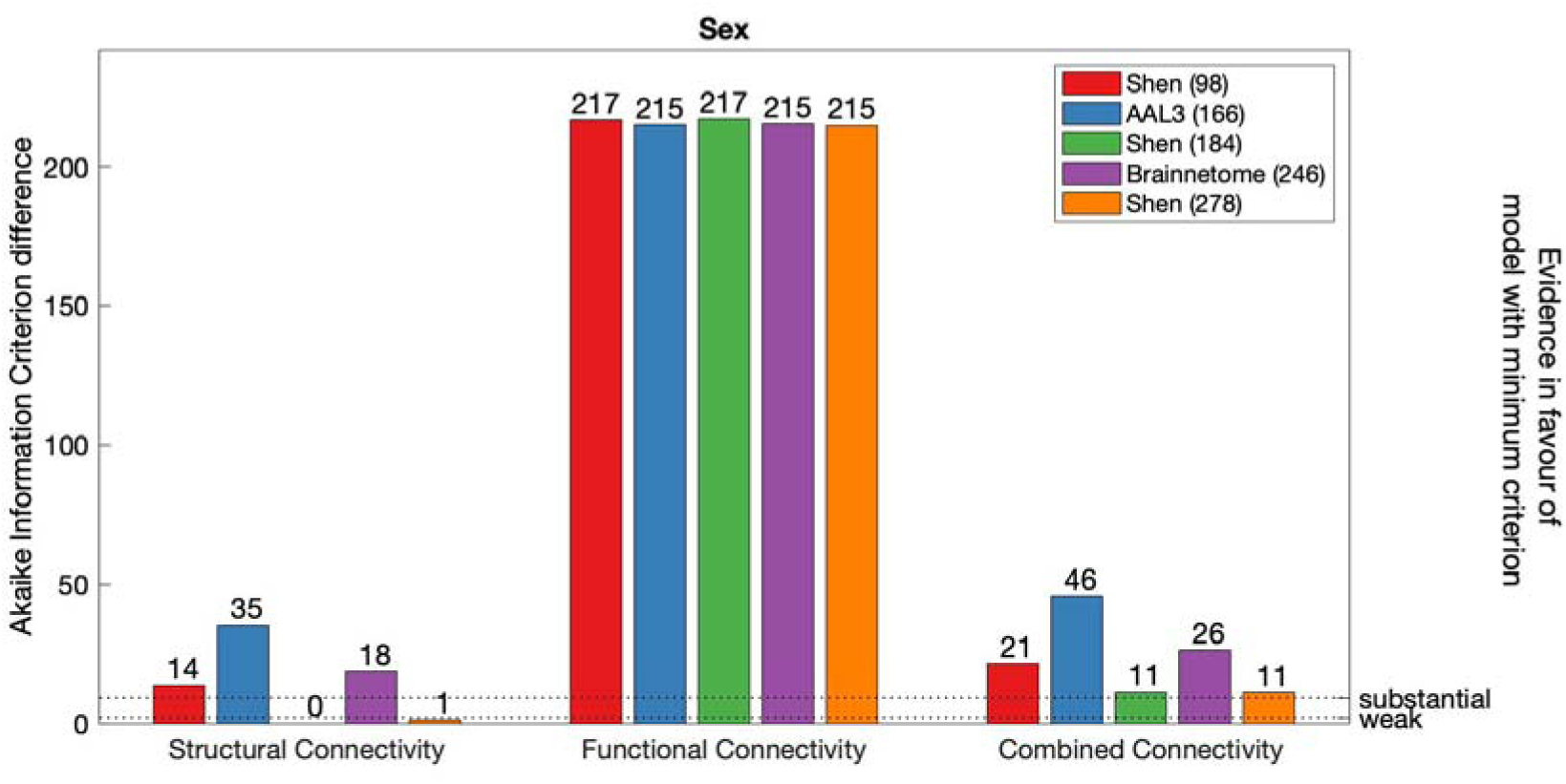
AIC difference for global graph theory measure models of Sex, constructed with each parcellation scheme. Dotted line marks weak, and substantial evidence in favour of SC model defined with Shen (184) (minimum AIC across all modalities and parcellations).

AIC values demonstrated that across all connectivity modalities and parcellations, age was best modelled with FC defined Brainnetome (246) parcellation (Figure 19). For SC, there was weak evidence in favour of Shen (93 and 184) and Brainnetome (246) relative to AAL3 (166). For FC, there was weak evidence in favour of Brainnetome (246) parcellation to AAL3 (166) and Shen (278) and strong-to-substantial evidence in favour of Brainnetome (246) to Shen (93 and 184). For CC, there was strong-to-substantial evidence in favour of Brainnetome (246) parcellation to all alternatives.

Next, across connectivity modalities and parcellations, education had lowest AIC value when modelled with SC defined with Brainnetome (246) or with FC defined with Shen (93 and 184) parcellation. Within SC, there was strong evidence in favour of Brainnetome (246) relative to alternative parcellations. For FC, there was no notable difference between AIC values of models defined with Shen (184) and Shen (93). For CC, there was weak-to-substantial evidence in favour of Brainnetome (246) relative to alternative parcellations.

Finally, there was substantial evidence in favour of SC and CC models of sex relative FC models. For SC and CC, lowest AIC values were estimated for SC models defined with either Shen (184) followed by Shen (278). These two models were substantially favoured to any other alternative models.

##### Model comparison based on out-of-sample performance

Results of the cross-validation of models of demographics are presented in Figures 22-24. It was found that graph theory measures of global organisation of SC, FC and CC could not produce out-of-sample predictions of demographics that would explain more variance in unseen samples than chance.

**Figure 22.**
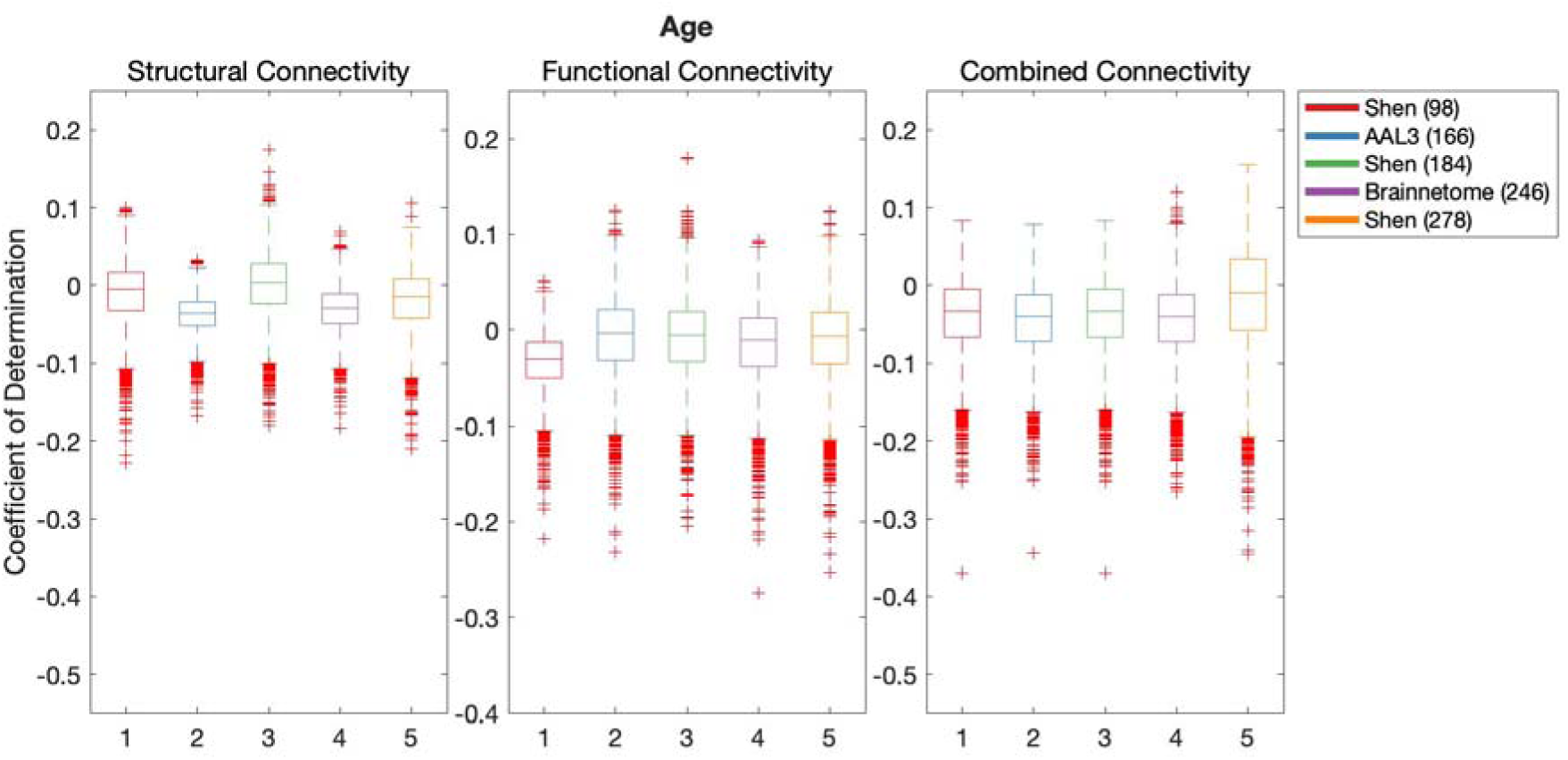
Results of BBC-CV of age models constructed with graph theory measures of SC, FC and CC, as measured by the coefficient of determination. The solid lines show the median scores, the boxes show the interquartile range (IQR), and ticks outside of whiskers indicate outlier scores across all bootstrap samples. Unfilled boxes illustrate bellow-chance prediction

**Figure 23.**
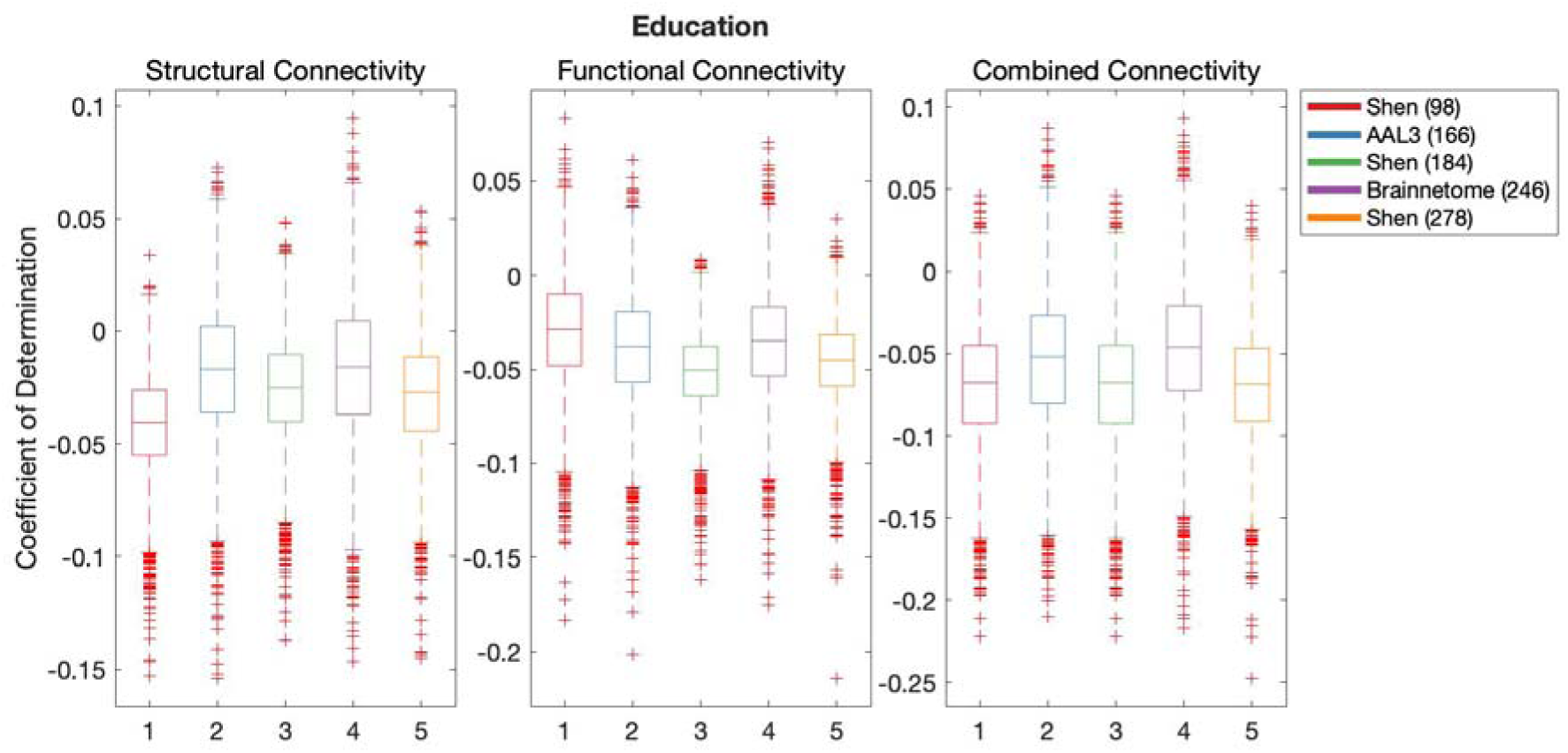
Results of BBC-CV of education models constructed with graph theory measures of SC, FC and CC, as measured by the coefficient of determination. The solid lines show the median scores, the boxes show the interquartile range (IQR), and ticks outside of whiskers indicate outlier scores across all bootstrap samples. Unfilled boxes illustrate bellow-chance prediction

**Figure 24.**
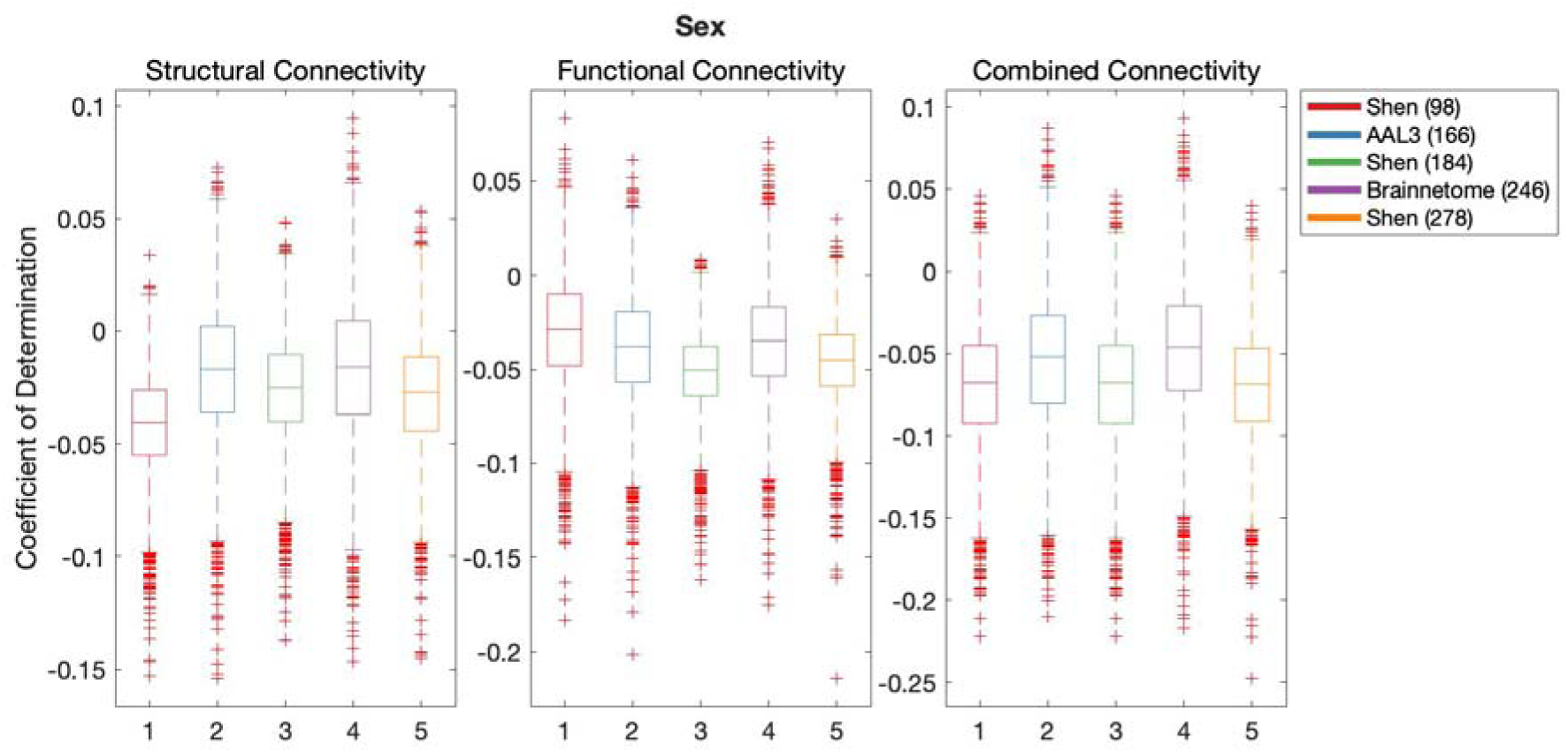
Results of BBC-CV of sex models constructed with graph theory measures of SC, FC and CC, as measured by the coefficient of determination. The solid lines show the median scores, the boxes show the interquartile range (IQR), and ticks outside of whiskers indicate outlier scores across all bootstrap samples. Unfilled boxes illustrate bellow-chance prediction

#### Predictive modelling of cognition

##### AIC-based model comparison

Figures 25-29 demonstrate AIC for models of cognition composed with global graph theory measures of SC, FC and CC when defined with the 5 parcellation schemes.

**Figure 25.**
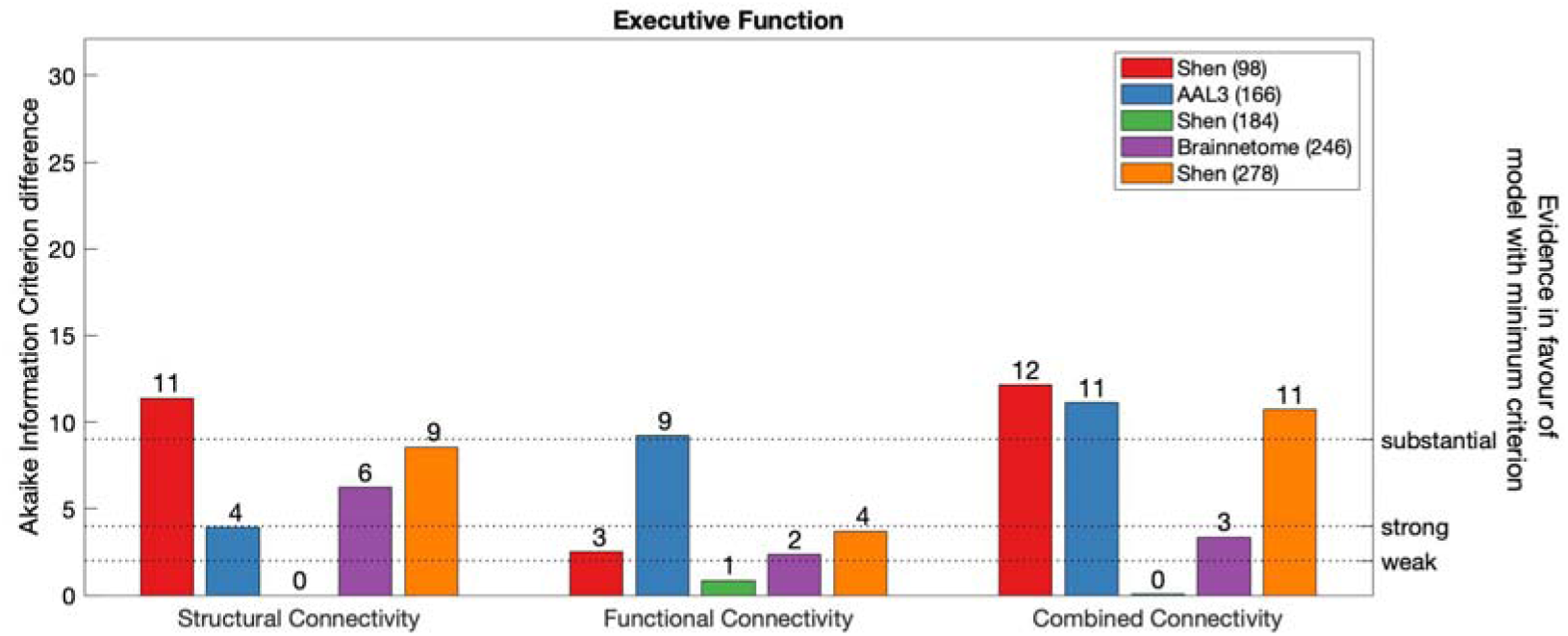
AIC difference for global graph theory measure models of Executive Function, constructed with each parcellation scheme. Dotted line marks weak, and substantial evidence in favour of SC model defined with Shen (184) (minimum AIC across all modalities and parcellations).

**Figure 26.**
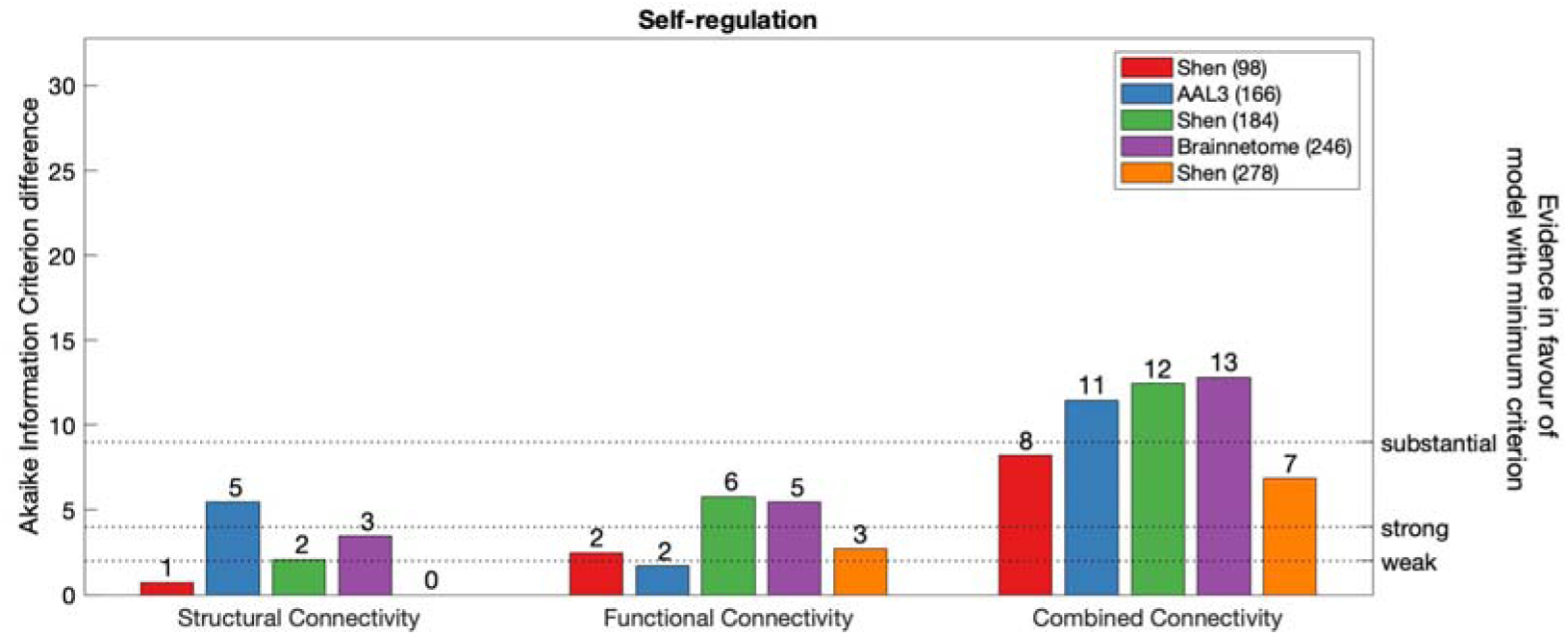
AIC difference for global graph theory measure models of Self-regulation, constructed with each parcellation scheme. Dotted line marks weak, and substantial evidence in favour of SC model defined with Shen (278) (minimum AIC across all modalities and parcellations).

**Figure 27.**
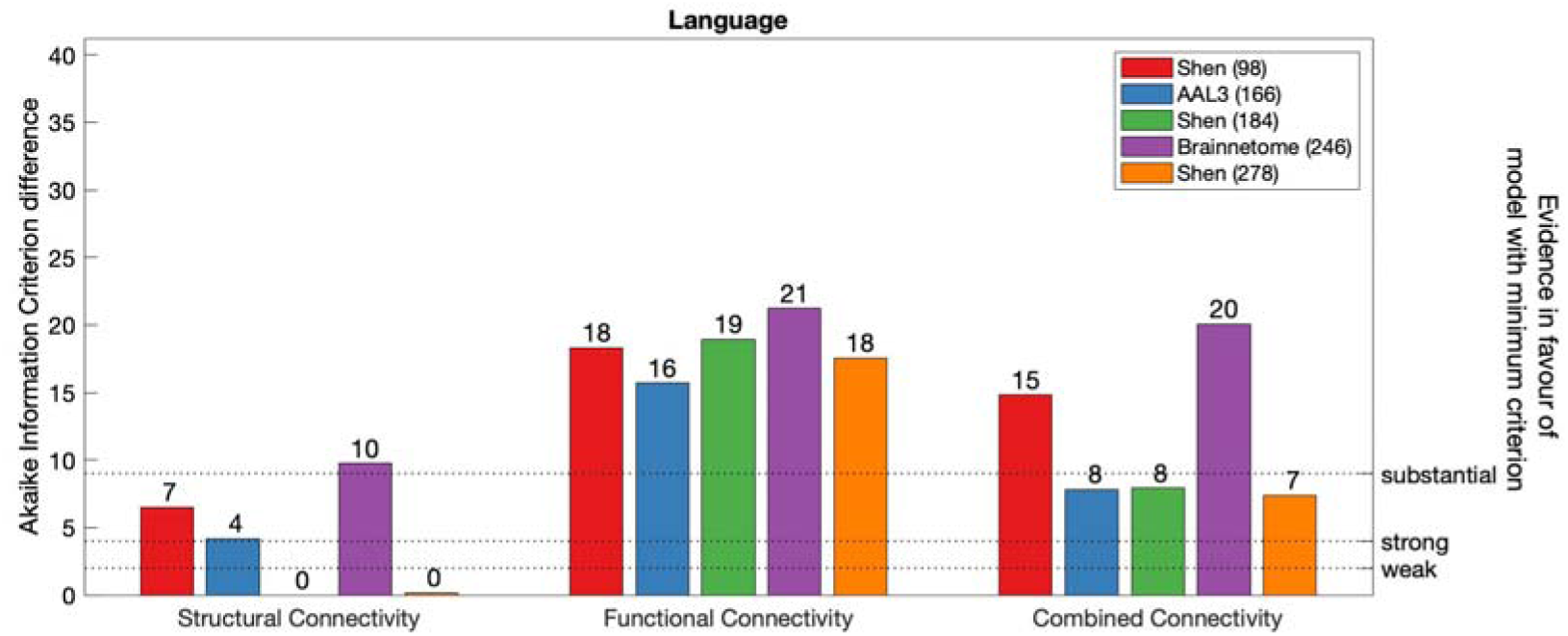
AIC difference for global graph theory measure models of Language, constructed with each parcellation scheme. Dotted line marks weak, and substantial evidence in favour of SC model defined with Shen (184) (minimum AIC across all modalities and parcellations).

**Figure 28.**
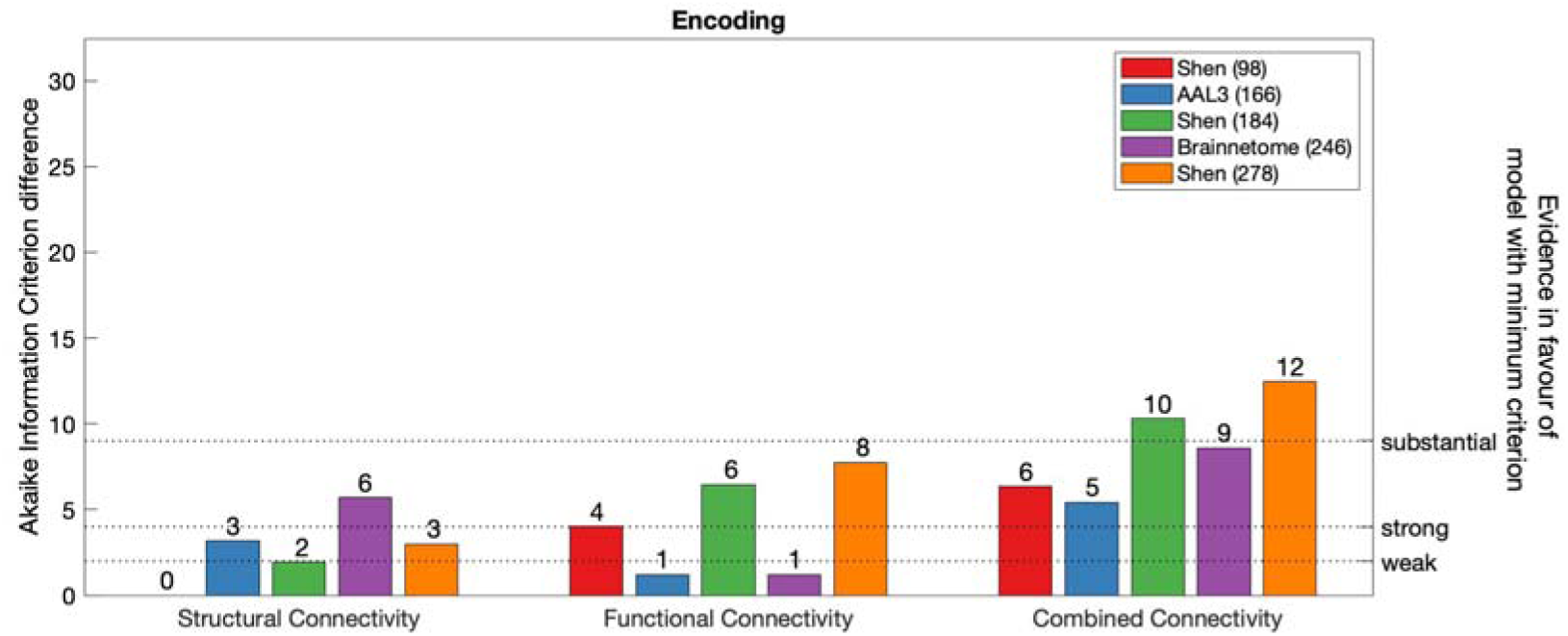
AIC difference for global graph theory measure models of Encoding, constructed with each parcellation scheme. Dotted line marks weak, and substantial evidence in favour of SC model defined with Shen (93) (minimum AIC across all modalities and parcellations).

**Figure 29.**
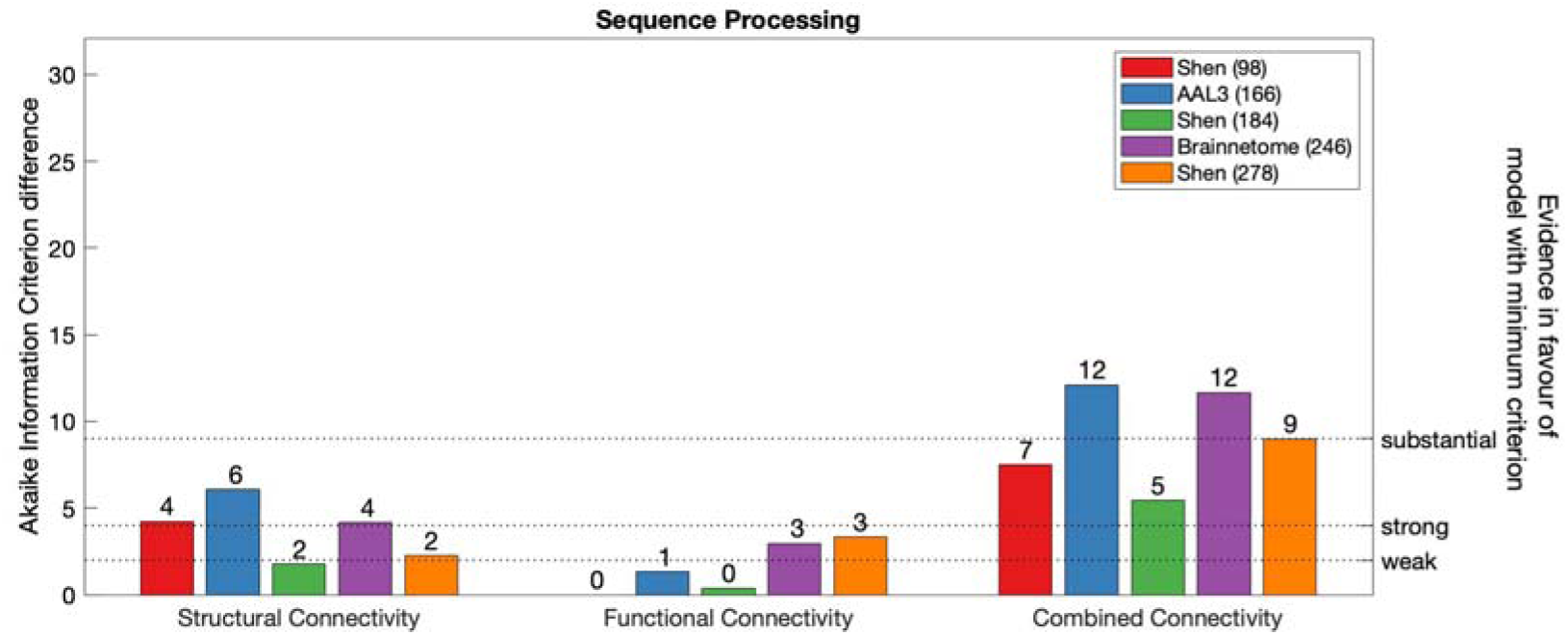
AIC difference for global graph theory measure models of Sequence Processing, constructed with each parcellation scheme. Dotted line marks weak, and substantial evidence in favour of SC model defined with Shen (93) (minimum AIC across all modalities and parcellations).

Overall, AIC favoured models of Executive Function composed with SC defined with Shen (184) parcellation. In addition, there was no notable difference between SC, FC and CC models of Executive Function defined with Shen (184) parcellation. For SC, FC and CC, there was strong-to-substantial evidence in favour of Shen (184) relative to alternative parcellations.

Self-regulation was best modelled by SC when parcellated with Shen (278). For SC, there was no notable difference between AIC values of Shen (278) and Shen (93). However, there was weak evidence in favour of Shen (278) relative to Shen (184) and Brainnetome (246), and strong evidence relative to AAL3 (166). For FC, Self-regulation was best modelled with AAL3 (166) but the evidence in favour of this model relative to Shen (93 and 278) was not notable, and it was weak relative to Shen (184) and Brainnetome (246). In For CC, there was no notable difference between AIC values of Shen (93) and Shen (278). However, there was strong evidence in favour of Shen (278) relative to AAL3 (166) and Shen (184), and substantial evidence relative to Brainnetome (246).

Language was best modelled by SC defined with Shen (184) parcellation. For SC, there was no notable difference between AIC values of Shen (184) and Shen (278), but there was strong to substantial evidence in favour of Shen (184) relative to alternative parcellations. For FC, AIC difference weakly favoured AAL3 (166) parcellation relative to Shen (93, 184), and strongly relative to Brainnetome (246). For CC, there was no notable difference between AIC values of Shen (278) and Shen (184) and AAL3 (166) but there was strong-to-substantial evidence in favour of these models relative to Shen (93) and Brainnetome (246).

Overall, Encoding was best modelled with SC defined with Shen (93) parcellation. When FC was used to model Encoding, best models were achieved with either AAL3 (166) or Brainnetome (246). When CC was used to model Encoding, according to AIC differences, the preferred model was defined with AAL3 (93) and Shen (93).

Finally, Sequence Processing was best modelled with FC defined with either Shen (93) or Shen (184). When either SC or CC was used, AIC values were in favour of models obtained with Shen (184).

##### Model comparison based on out-of-sample performance

Results of the cross-validation of models of cognition are presented in Figures 30-34. It was found that graph theory measures of global organisation of SC, FC and CC could not produce out-of-sample predictions of cognition that would explain more variance in unseen samples than chance.

**Figure 30.**
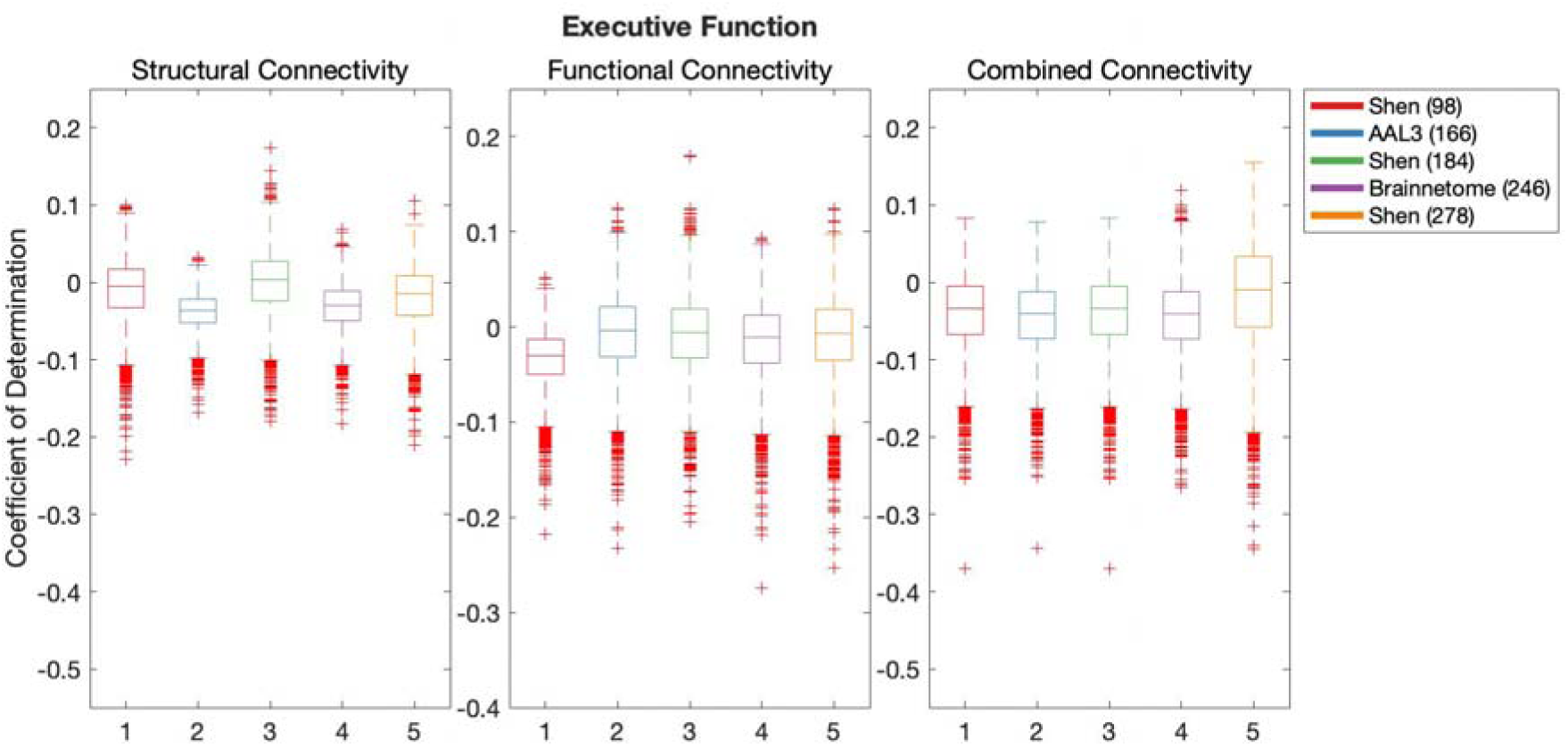
Results of BBC-CV of Executive Function models constructed with graph theory measures of SC, FC and CC, as measured by the coefficient of determination. The solid lines show the median scores, the boxes show the interquartile range (IQR), and ticks outside of whiskers indicate outlier scores across all bootstrap samples. Unfilled boxes illustrate bellow-chance prediction

**Figure 31.**
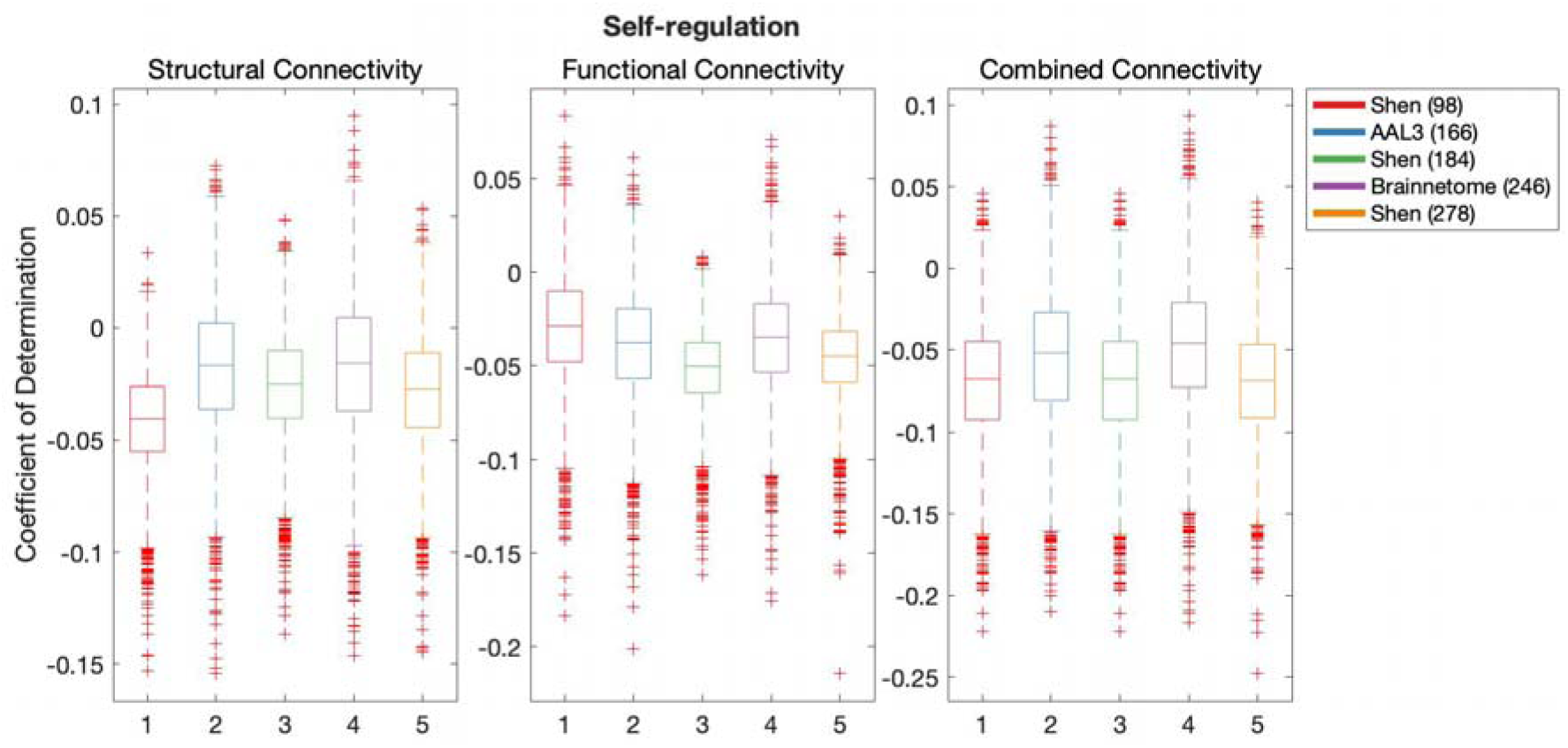
Results of BBC-CV of Self-regulation models constructed with graph theory measures of SC, FC and CC, as measured by the coefficient of determination. The solid lines show the median scores, the boxes show the interquartile range (IQR), and ticks outside of whiskers indicate outlier scores across all bootstrap samples. Unfilled boxes illustrate bellow-chance prediction

**Figure 32.**
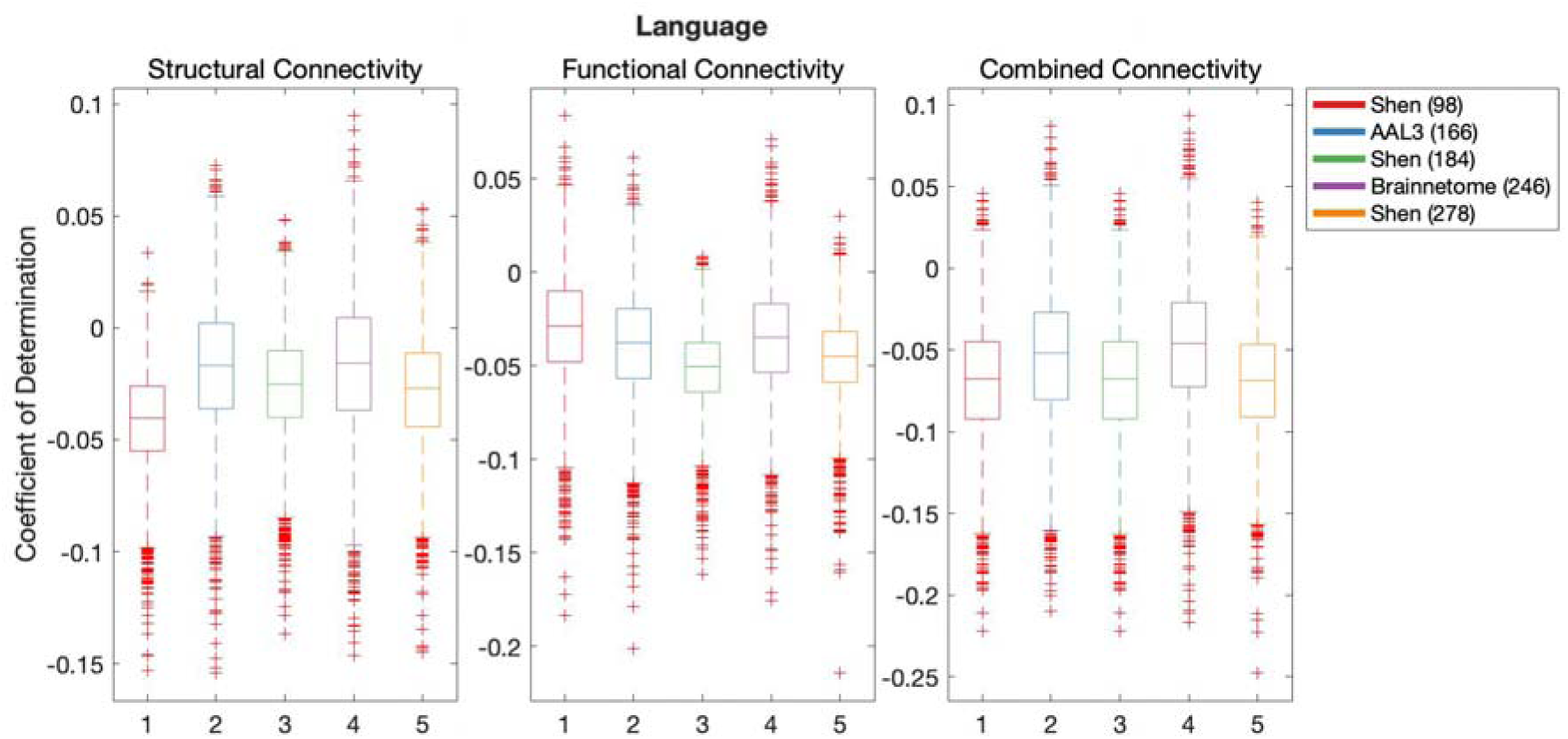
Results of BBC-CV of Language models constructed with graph theory measures of SC, FC and CC, as measured by the coefficient of determination. The solid lines show the median scores, the boxes show the interquartile range (IQR), and ticks outside of whiskers indicate outlier scores across all bootstrap samples. Unfilled boxes illustrate bellow-chance prediction

**Figure 33.**
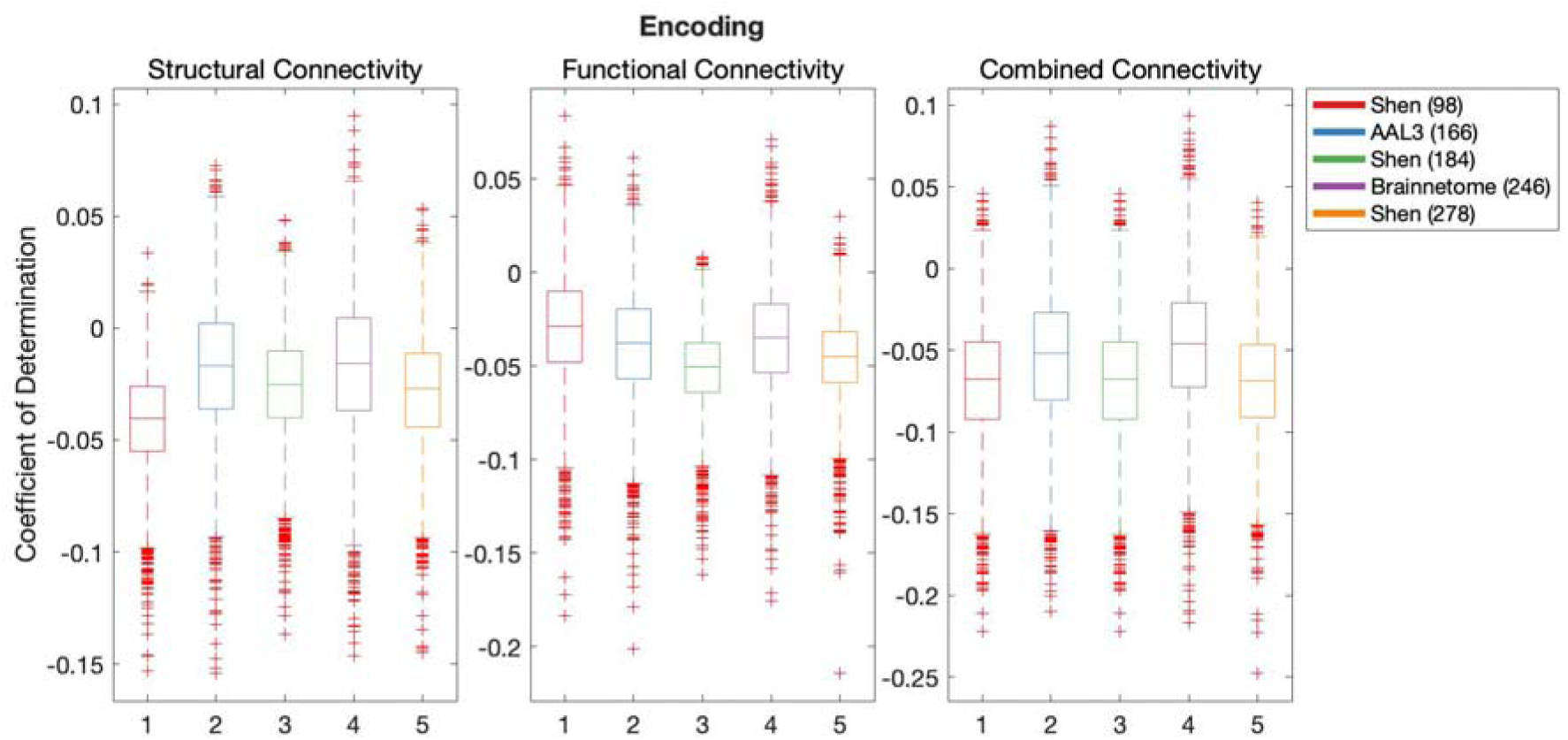
Results of BBC-CV of Encoding models constructed with graph theory measures of SC, FC and CC, as measured by the coefficient of determination. The solid lines show the median scores, the boxes show the interquartile range (IQR), and ticks outside of whiskers indicate outlier scores across all bootstrap samples. Unfilled boxes illustrate bellow-chance prediction

**Figure 34.**
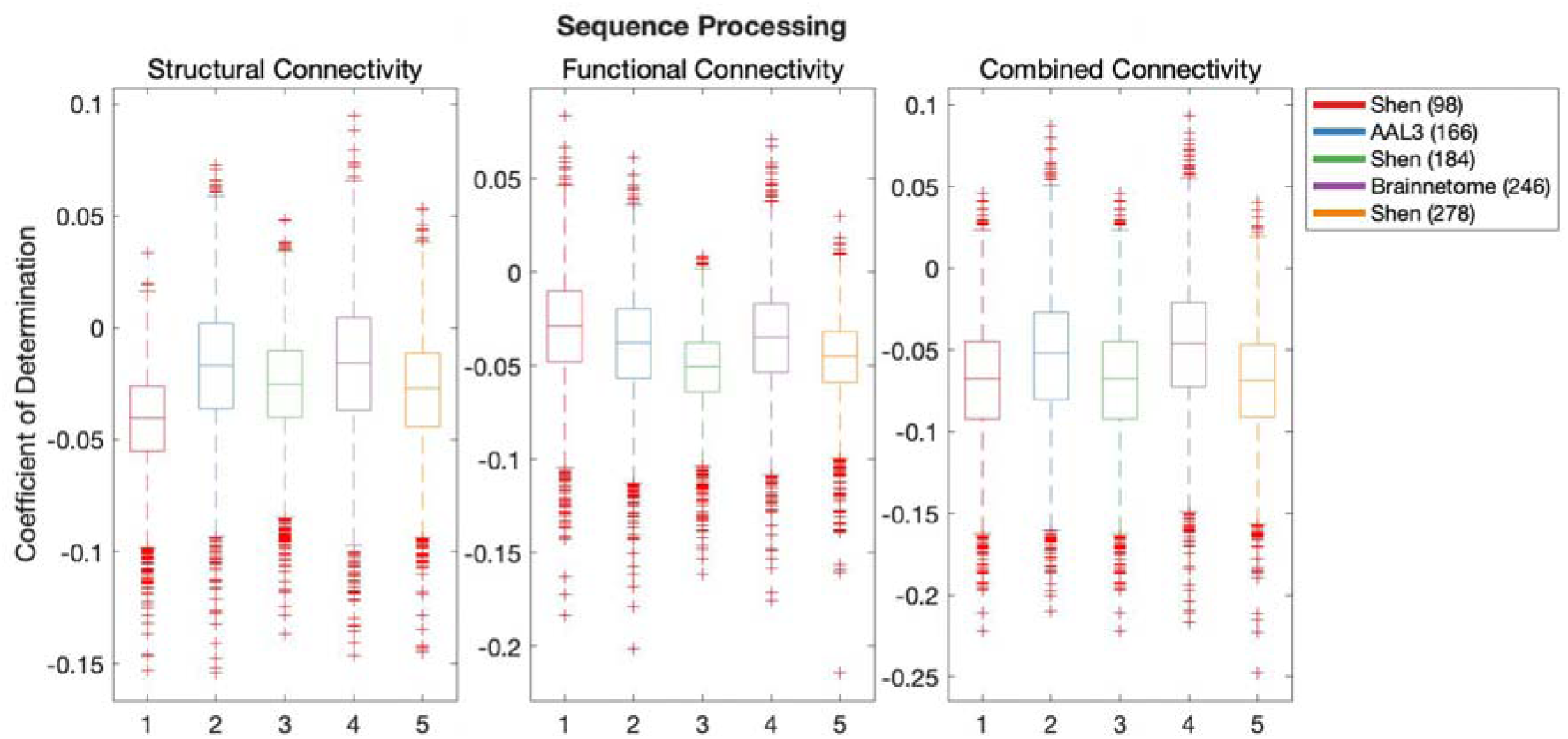
Results of BBC-CV of Sequence Processing models constructed with graph theory measures of SC, FC and CC, as measured by the coefficient of determination. The solid lines show the median scores, the boxes show the interquartile range (IQR), and ticks outside of whiskers indicate outlier scores across all bootstrap samples. Unfilled boxes illustrate bellow-chance prediction

## DISCUSSION

Currently available brain parcellation schemes vary in the neuroimaging data and the node-generation algorithms used to obtain them. Consequently, the obtained parcellations vary in the number, size and shape of nodes. These differences have been demonstrated to largely impact the predictive power of connectivity-based models of cognition (Dhamala et al., 2021; Mellema et al., 2020; Ota et al., 2014; Pervaiz et al., 2020). However, to date, it remained unclear if the method of parcellation is systematically impacting predictive models. To fill this gap in literature, the present research defined SC, FC and CC with 5 different popular parcellation schemes and constructed predictive models of demographics and cognition based on models using each connectivity modality and parcellation scheme as predictors. Results of AIC-based model comparison did not show notable benefit to choosing a specific parcellation to model sample demographics. However, there was some trend towards the benefit of using the low-resolution functional parcellation (Shen 93) to model cognition. Analysis of the out-of-sample performance of the models showed that most parcellations produced above-chance predictive performance of demographics for all connectivity modalities. The only exception to this was the model of educational level using FC with the Shen 278 parcellation. In addition, it was found that models of demographics showed higher performance when using FC defined with functional parcellations. There was no consistent benefit to using low or high-resolution parcellations for this connectivity modality. When SC was used to model age and education, AAL3 (166) parcellation produced the highest out-of-sample performance. In contrast, models of sex based on SC modality benefitted from using low-resolution functional parcellation (Shen, 93). When out-of-sample predictions of cognition were assessed, it was found that no parcellation scheme would consistently produce significantly higher predictive performance than the others, even when the connectivity and parcellation corresponded to the same modality (functional or structural). However, out-of-sample predictions of cognition were consistently above chance when using connectivity defined with the Shen 93 parcellation scheme. This was the only parcellation scheme that achieved such consistency in performance.

Prior research has demonstrated that effective modelling of cognition to some extent depends on parcellation. For example, Dhamala et al. (2021) reported that SC defined with low-resolution FreeSurfer parcellation succeeded in producing predictions of crystallized intelligence in an unseen sample. However, the same connectivity failed to produce effective predictions when in-house high-resolution parcellation (CoCo 439) was used instead. In contrast, FC tended to benefit at predicting cognitive abilities when CoCo 439 parcellation was used. Their CoCo 439 parcellation was defined from functional data suggesting a benefit of using a parcellation that corresponds to the modality of connectivity. However, in the present report, when individual cognitive domains were analysed such benefits were lost. For example, we found that Executive Function was most effectively modelled with FC (as reflected by out-of-sample predictions), when FC was defined with AAL3 (166) parcellation – a parcellation defined with neural anatomy. In addition, similarly to the report from Dhamala et al. (2021), this work could not identify a single superior parcellation. Such inconsistency of findings warrants caution when interpreting results obtained with predictive modelling. Authors must consider that any superiority of a given model defined with a given set of data may be related to the parcellation scheme used.

In another seminal report, Pervaiz et al. (2020) demonstrated that the most accurate predictions of fluid cognition were produced with models defined with high-resolution parcels generated by spatial Independent Component Analysis. With this algorithm, parcels can overlap and are not necessarily contiguous. This contrasts with publicly available parcellations used in the present work, which were defined to identify non-overlapping, contiguous parcels. Consequently, the parcellations offered with fewer constraints may prove to produce more accurate parcels with coherent signals (i.e. avoid partial voluming of signals). For example, recent work has demonstrated graded changes in connectivity patterns across cortex (Bajada et al., 2017; Cloutman et al., 2020), which suggests that smooth parcel boundaries may better describe connectivity of the system. This need for overlapping parcels may explain why the results presented by Pervaiz et al. (2020) identified high resolution to benefit modelling of fluid cognition, while our results could not identify this pattern. However, Pervaiz and colleagues (2020) have not assessed the predictive value of various parcellation schemes in modelling done with different parcellations with SC or CC. The present work adds to the field the finding that SC and CC do not benefit in predictive power from modality-corresponding parcellation or a specific resolution of parcellation.

One consistent finding across models of demographics and cognition was that the low-resolution functional parcellation (Shen, 93) always produced predictions that explained more variation than chance. This was observed regardless of whether SC, FC or CC was used to predict demographics and cognition. One explanation of this finding is that higher-resolution parcellations may be more prone to partial voluming of signals due to individual differences in neural anatomy and function. This would result in models that are more specific to the training sample and decreased generalisability skills. However, where higher-resolution parcellation schemes succeeded at modelling demographics and cognition, they always proved to produce more generalisable models. This highlights that both low-resolution parcellations may offer some insights into neural substrates of cognition and high-resolution further complement this information.

A key finding of this work was that although parcellation schemes impacted the global architecture of brain networks, the global architecture of the networks had no out-of-sample predictive power with demographics or cognition. This suggests that global brain organisation measures reflect changes in the organisation across parcellation schemes but this change alone cannot account for the reason why specific parcellation schemes excel at modelling demographics or cognition. At the moment, partial voluming of signals presents a much stronger explanation of why specific parcellations succeed at modelling cognition and others fail. In the future, it will be possible to implement in-house parcellations that allow for overlapping parcels and manipulate the resolution of parcellation (e.g. spatial Independent Component Analysis, spectral clustering and spectral-reordering algorithm), to generate different parcellations and investigate further if organisational properties revealed by specific resolution of parcellation relate to demographics and cognition.

One key limitation of the present work is that we have not considered alternative resolutions of parcellations defined with anatomical or a hybrid of anatomical and functional information. In particular, we have investigated 3 resolutions of functional parcellation but only a single resolution of anatomical and hybrid parcellation. It is possible that investigations of alternative resolutions in these corresponding modalities of connectivity would reveal a distinct benefit for predictive modelling with SC or CC. For example, it is possible that high- or low-resolution SC is particularly important for the prediction of cognitive domains and we failed to capture this because we only varied resolution for the functional parcellation scheme. A functional parcellation scheme is likely poorly suited to accurately identify parcel volumes in structural connectivity and it may have missed the benefit of various resolutions of structural parcellations.

Overall, this paper has investigated whether the method of parcellation is systematically related to the efficiency of predictive models of cognition and parcellation. By investigating the generalisability of SC, FC and CC predictive models defined with 5 different parcellations, we have found that low-resolution parcellation (Shen 93) was consistently able to produce generalisable models of demographics and cognition. However, when alternative parcellations succeeded at producing generalisable models then these parcellations outperformed the Shen 93 parcellation. It remained challenging to identify a superior parcellation scheme. As a result of this work, it is important that from here onwards authors provide a cautious interpretation of superior models of cognition based on connectivity, as specific parcellation schemes may unsystematically impact the results.

